# Assessing the Performance of qpAdm: A Statistical Tool for Studying Population Admixture

**DOI:** 10.1101/2020.04.09.032664

**Authors:** Éadaoin Harney, Nick Patterson, David Reich, John Wakeley

## Abstract

qpAdm is a statistical tool for studying the ancestry of populations with histories that involve admixture between two or more source populations. Using qpAdm, it is possible to identify plausible models of admixture that fit the population history of a group of interest and to calculate the relative proportion of ancestry that can be ascribed to each source population in the model. Although qpAdm is widely used in studies of population history of human (and non-human) groups, relatively little has been done to assess its performance. We performed a simulation study to assess the behavior of qpAdm under various scenarios in order to identify areas of potential weakness and establish recommended best practices for use. We find that qpAdm is a robust tool that yields accurate results in many cases, including when data coverage is low, there are high rates of missing data or ancient DNA damage, or when diploid calls cannot be made. However, we caution against co-analyzing ancient and present-day data, the inclusion of an extremely large number of reference populations in a single model, and analyzing population histories involving extended periods of gene flow. We provide a user guide suggesting best practices for the use of qpAdm.

## INTRODUCTION

The last decade has experienced a revolution in the amount of genetic data available to study from both living and ancient organisms. Questions about the origins of populations have increased in complexity, often in an effort to understand histories that involve admixture, which are incompatible with traditional tree-like models of relatedness. qpAdm is a tool that can be used to understand the history of admixed populations. It has been applied to study the genetic history of human populations that would otherwise remain mysterious. For instance, the use of qpAdm was vital to studying the ancestry of the Late Bronze Age Greek culture of the “Mycenaeans” (Lazaridis *et al.* 2017)—the subjects of the Iliad and Odyssey. However, little has been done to assess qpAdm’s performance under both simple and complex scenarios.

A potential drawback of many population genetic tools for studying the population history of specific groups (Patterson *et al.* 2012; Pickrell and Pritchard 2012) is that they require the historical relationships of all other populations included in the analysis to be explicitly modeled. This underlying phylogeny is either specified by the user (as in qpGraph) or is calculated during the analysis (as in TreeMix). This may lead to biases or errors in inferences about admixture if mistakes are made when specifying the underlying relationships of non-target populations (Lipson 2020). This requirement for a complete and accurate population history is especially difficult to satisfy in studies that utilize ancient DNA, which increasingly attempt to use genetic data of limited quality to analyze nuanced differences between closely related groups. However, even in cases where it is difficult to reconstruct a full population history, it is often possible to examine patterns of shared genetic drift between various populations in order to learn about their relationship relative to one another (Patterson *et al.* 2012). qpAdm exploits this information, enabling admixture models to be tested for plausibility and admixture proportions to be estimated.

The theory underlying qpAdm, which was introduced in Haak *et al.* (2015), builds upon a class of statistics known as *f*-statistics (Patterson *et al.* 2012). *f*-statistics analyze patterns of allele frequency correlations among populations in order to determine whether their population histories can be described using strictly tree-based models, or if more complex models, such as those involving admixture, are required to explain the genetic data. *f*-statistics have been widely used in the population genetic literature and their behavior is well understood (Reich *et al.* 2009; Patterson *et al.* 2012; Peter 2016; Soraggi and Wiuf 2019; Lipson 2020). qpAdm harnesses the power of *f*-statistics to determine whether a population of interest (a target population) can be plausibly modeled as descending from a common ancestor of one or more source populations. For example, in a model with two source populations, qpAdm tests whether the target population is the product of a two-way admixture event between these source populations. The method requires a list of target and source populations and a list of additional reference populations which provide information about the relationships among the target and source populations.

The target and source populations are collectively referred to as ‘left’ populations, due to their position as the left-most arguments in the *f*_4_-statistics involved in the calculations. Additionally, users must specify a set of ‘right’ populations that serve as references against which the relationships of the target and source populations are considered. Previously, ‘right’ populations were referred to as ‘outgroup’ populations, but we avoid this term because it suggests that reference populations should be outgroups in phylogenetic sense (i.e. equally closely related to all ‘left’ populations). In fact, if all ‘right’ populations are symmetrically related to all ‘left’ populations in this way, qpAdm will not produce meaningful results. The method requires differential relatedness, meaning that at least some ‘right’ populations must be more closely related to a subset of ‘left’ populations than to the other ‘left’ populations. We illustrate this further in Methods & Results.

qpAdm computes a matrix of *f*-statistics of all possible pairs of populations in the ‘left’ and ‘right’ sets, of the form *f*_4_(Left_i_, Left_j_; Right_k_, Right_l_). If the source populations are descended from n different ancestral populations, then the matrix of *f*-statistics will have a rank (a maximum number of linearly independent allele-frequency vectors) equal to n-1 (Haak *et al.* 2015). We note that if all *f*_4_-statistics are computed from the same set of SNPs, which is the default mode of qpAdm, a basis for the statistics can be found using a matrix of reduced dimension, specifically by fixing both the target population (Left_*i*_ above) and Right_*k*_. This also improves the efficiency of the covariance calculations. qpAdm accounts for correlations between neighboring alleles and between related populations, measuring covariance using a block jackknife. For each model, it gives a p-value which is used to determine whether the proposed admixture model is plausible. The p-value is calculated using a likelihood ratio test comparing a constrained null model to an unconstrained alternative model. Specifically, it tests whether including the target population in the ‘left’ populations requires an additional independent ancestral population (i.e. changes the rank of the matrix of *f*-statistics from n-1 to n). A simple example in which this would be required, and the constrained model would be rejected, is when the putative target population is actually an outgroup to all source populations. In the constrained model, the admixed ancestral population of the target population is a mixture of sources.

While qpAdm has been theoretically described (Haak *et al.* 2015) and applied in numerous studies (e.g. Lazaridis *et al.* 2016; Haber *et al.* 2017; Lazaridis *et al.* 2017; Skoglund *et al.* 2017; de Barros Damgaard *et al.* 2018a; de Barros Damgaard *et al.* 2018b; Hajdinjak *et al.* 2018; Harney *et al.* 2018; Narasimhan *et al.* 2018; Olalde *et al.* 2018), producing results that are consistent with those of other population genetic methods, very little has been done to assess the performance of the tool when the population history is known (i.e. using simulated data). The only simulation-based analysis that has been previously conducted examined whether simulated populations—generated according to the model fitted by qpAdm, by resampling data using the source populations and estimated admixture proportions—behaved similarly to the real target population in further statistical analyses (Lazaridis *et al.* 2017). Although this limited example supports the use of qpAdm in population genetic analyses, it did not address any of the potential limitations of the method. Here we use simulated genomic data to study the distributions of p-values and estimated admixture proportions from qpAdm, the potential of qpAdm to distinguish optimal from non-optimal models of admixture for a given set of samples, and the performance of qpAdm in the face of more challenging demographic scenarios.

The chief purpose of qpAdm is to identify a subset of plausible models of a population’s ancestry from a larger set of possible models. Models are deemed implausible if they are rejected (by having a small p-value) or if their estimated admixture proportions fall outside the biologically relevant range (0,1). Thus, p-values are applied in a non-standard statistical way. Users propose a range of possible models, in which they attempt to model the target population using a variety of different combinations of populations as sources, then eliminate implausible models. The set of plausible models are the ones which are *not* rejected, meaning they have p-values *greater* than the chosen significance level, which is usually 5%. To illustrate, Box 1 describes how an analogous technique might be applied to identify plausible values of the (unknown) probability of heads for a coin.

Identical to standard statistical methods, this approach will work best when the p-values generated by qpAdm follow a uniform distribution, if the correct admixture model is specified. Then the correct model will be rejected 5% of the time when a threshold of p<0.05 is applied. For other plausible but less-optimal models, the distribution of p-value is not expected to be uniform but should have an appreciable chance of being above the 5% cutoff. The distribution of p-values for implausible or incorrect models should fall largely below the 5% cutoff. While experience suggests that the p-values generated by qpAdm are reasonably consistent with these expectations, in this work we perform the first systematic test of these ideas.

### Box 1.

#### Coin flipping analogy

Imagine that we wish to know which of several possible models (here values, p_0_) of the probability of heads best describes the behavior of a coin. The actual value is unknown and the coin may be unfair. To mimic what is done in qpAdm, we might specify a set of possible models, for instance with probabilities of heads equal to 0.1, 0.2, 0.3, 0.4, 0.5, 0.6, 0.7, 0.8, or 0.9. To determine which of the possible models are plausible for the coin, we flip it multiple times and count the number of heads we observe. The probability of observing that number of heads under each of the possible models would be given by the binomial distribution. Again by analogy with qpAdm, we could assess the plausibility of each model by a generalized likelihood ratio test of each null model against the unconstrained alternative.

If we flip the coin 10 times, and it lands on heads 7 times, using a p-value threshold of 0.05, we can already eliminate 0.1-0.4 and 0.9 as potential models of the probability of heads for our coin (Table 1.1). By increasing the number of flips to 100, if we observe 73 heads, we can further eliminate 0.5 and 0.6, leaving only 0.7 and 0.8 as plausible models for the probability of heads of the coin. As with this analogy, qpAdm identifies a set of plausible models by not rejecting null hypotheses (possibly by what would be Type II error in a standard statistical test).

Note that in this coin flipping analogy, any model which did not have the exactly correct probability of heads would eventually be rejected if enough data were collected. Our findings in the main text suggest that this particular problem is not an issue for qpAdm, which specifies models in a different way and which admits a range of similar, plausible models even using whole-genome data.

**Table 1.1.** Using a generalized likelihood-ratio test of H_0_:p=p_0_ to identify plausible models for the probability of heads of a coin. Models that produce p-values greater or equal to 0.05 are highlighted in blue, while those that are less than 0.05 are highlighted in red.

| Models ($p_0$) for the probability of heads | P-values from a generalized likelihood ratio test of $H_0:p=p_0$ against the unconstrained alternative | |
| --- | --- | --- |
| Number of flips | 10 | 100 |
| Number of heads | 7 | 73 |
| 0.1 | < 0.001 | < 0.001 |
| 0.2 | < 0.001 | < 0.001 |
| 0.3 | < 0.001 | < 0.001 |
| 0.4 | 0.007 | < 0.001 |
| 0.5 | 0.070 | < 0.001 |
| 0.6 | 0.353 | 0.006 |
| 0.7 | 1.000 | 0.508 |
| 0.8 | 0.289 | 0.092 |
| 0.9 | 0.013 | < 0.001 |

Similarly, although the estimated admixture proportions calculated by qpAdm appear generally consistent with values generated using other statistics, the accuracy of these estimates have never been rigorously tested. Of particular interest is the accuracy of these estimates when calculated on low quality data, as qpAdm is often applied to the study of ancient DNA, which is characteristically low coverage, may have a high rate of missing data, and is susceptible to deamination of cystosine nucleotides (manifesting in sequence data as cytosines being misread as thymines). Further, ancient DNA is often subject to a complex ascertainment process that could potentially bias statistical analyses. We explore the impact of each of these factors on the admixture proportions estimated by qpAdm.

Additionally, while one of the main features of qpAdm is its ability to distinguish between optimal and non-optimal models for a group’s population history, there are no formal recommendations about what strategy should be employed to compare models. We therefore consider two of the most commonly employed strategies for model comparison, highlighting their potential benefits and weaknesses.

Finally, we conclude by exploring non-standard cases where the expected behavior of qpAdm is poorly understood, such as the impact of including a large number of populations in the reference population set and the behavior of qpAdm when applied to population histories that involve continuous gene flow rather than single pulses of admixture.

We show that qpAdm reliably identifies population histories involving admixture and accurately infers admixture proportions. It is robust to low coverage, high rates of missing data, DNA damage (when occurring at similar rates in all populations), the use of pseudo-haploid data, small sample size, and ascertainment bias. We also identify some issues with naive applications of qpAdm. One of these issues is that multiple plausible scenarios may be found most of which are not the truth because qpAdm uses non-rejection of null models as its criterion for plausibility. Another of these issues is that true models may be rejected if samples from too many populations are included in the analysis. A third is that qpAdm results may be difficult to interpret and even misleading under conditions of continuous gene flow. In order to help guard against these potential pitfalls and make this tool more accessible to users, we include an updated user guide for qpAdm (Supplementary Materials 1) and make specific recommendations for best practices for use.

## METHODS & RESULTS

### Data Generation

We used msprime version 0.7.1 (Kelleher *et al.* 2016) to simulate genome-wide data using the TreeSequence.variants() method, which provides information about all mutations arising in the dataset and the genotype of individuals at variant sites. We then converted this output to EIGENSTRAT format (Patterson *et al.* 2006). Parameters were chosen in order to mirror what has been estimated for humans, including a mutation rate of 1.5×10^−8^ mutations per base pair per generation, recombination rate of 1.0×10^−8^ per base pair per generation, and effective population sizes between 2.5×10^4^ and 8.0×10^5^ (varying between populations and over time; see Supplementary File 1 for full details). We generated sequence data for 22 chromosomes, each of the approximate length of each of the human autosomes. We simulated 2*n haploid individuals then combined pairs of haploid individuals to form n diploid individuals.

In order to assess the performance of qpAdm when the population history of a group is relatively simple and fully understood, we simulated genetic data according to a base population tree (Figure 1), consisting of 16 populations and two admixture events (one relatively recent and the other occurring much earlier in the population history). For the more recent admixture event, lineages 14a and 14b contribute α and 1− α proportion of ancestry to population 14, respectively. Unless otherwise noted, α is equal to 0.5. In the earlier admixture event, lineages 15a and 15b contribute β and 1-β proportion of ancestry to population 15, respectively, where β is equal to 0.55. This tree is an expanded version of a population tree described in Patterson *et al.* (2012), which was used to test the performance of the tool qpGraph. The exact simulation parameters we used are described in Supplementary File 1. These were chosen so that the overall level of variation (total number of SNPs) and the differentiation between populations (F_ST_) were similar to what is observed for humans.

**Figure 1.**
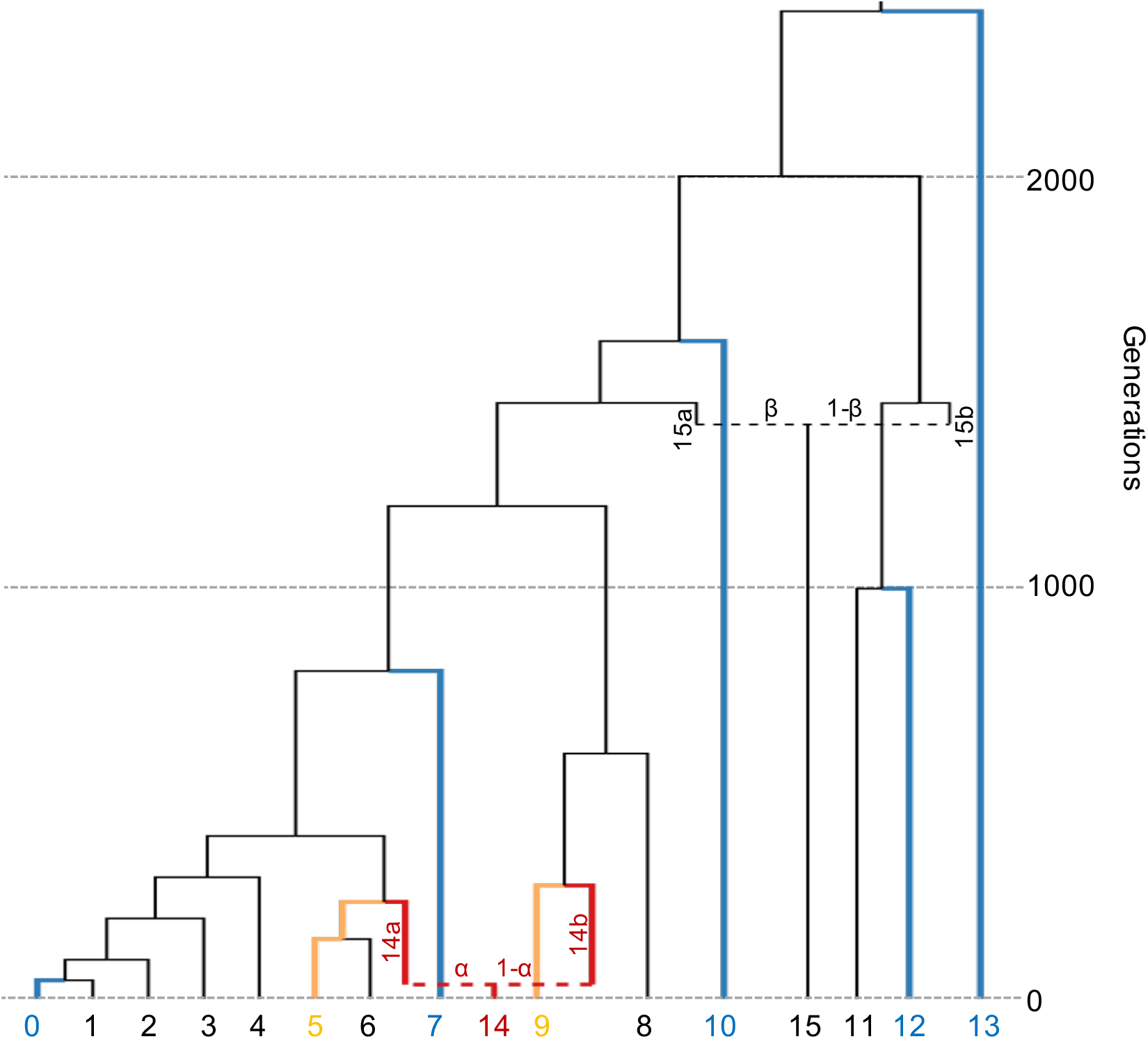
Population history of simulated data. Populations included in the standard model used for qpAdm models are indicated as follows: target (red), sources (yellow), references (blue).

For most of our simulations, we generated genomic data for samples taken from 10 (diploid) individuals from each of the 16 populations in Figure 1. The populations in Figure 1 are idealized, theoretical populations (see Winther *et al.* 2015) and are not meant to represent any particular human groups. Likewise, the mostly tree-like relationships of populations in Figure 1 simply reflect the kinds of historical scenarios qpAdm was designed to handle. We consider an example of non-tree-like structure in the section on continuous gene flow.

Unless otherwise noted, the admixture model of interest is defined as follows; population 14 is the target population (the ancestry of which is being modeled), populations 5 and 9 are defined as the sources of this admixture, while populations 0, 7, 10, 12, and 13 are designated as reference populations. As none of these reference populations are more closely related to the target population than to either of the two source populations (i.e. the reference populations do not have any shared drift with the target population that is not also shared with at least one of the source populations), this model should be considered plausible. This model will be referred to as the standard model. Note that because populations 5 and 6 are symmetrically related to population 14, both represent be equally good sources of its ancestry. Unless otherwise noted, population 6 will therefore be excluded from analyses.

All qpAdm analyses were performed using qpAdm version 960, using default parameters, and the optional parameters, “allsnps: YES”, “details: YES” and “summary: YES”, unless otherwise specified. See Supplementary Materials 1 for a complete description of all qpAdm parameters.

### Distribution of p-values

qpAdm outputs a p-value that is used to determine whether a specific model of population history can be considered plausible. Models are rejected, or regarded as implausible, when the p-value is below the chosen significance cutoff (typically 0.05). In order for true models to be rejected properly at this nominal significance level, that is only 5% of the time, the distribution of p-values should be uniform when the null model is equal to the true model. However, this assumption of uniformity of p-values in qpAdm has never been confirmed. We therefore assessed the distribution of p-values produced by qpAdm by simulating 5,000 replicates under our standard model (defined in Figure 1) and running qpAdm on each replicate using the target, source and reference populations defined in the standard model. We find that the p-values generated by qpAdm appear uniformly distributed (Figure 2A; Supplementary Table 1). Using a Kolmogorov-Smirnov test, we fail to reject the null hypothesis that the calculated p-values are uniformly distributed (p=0.644), supporting theoretical predictions for the uniform distribution of p-values generated by qpAdm when an accurate model is presented.

**Figure 2.**
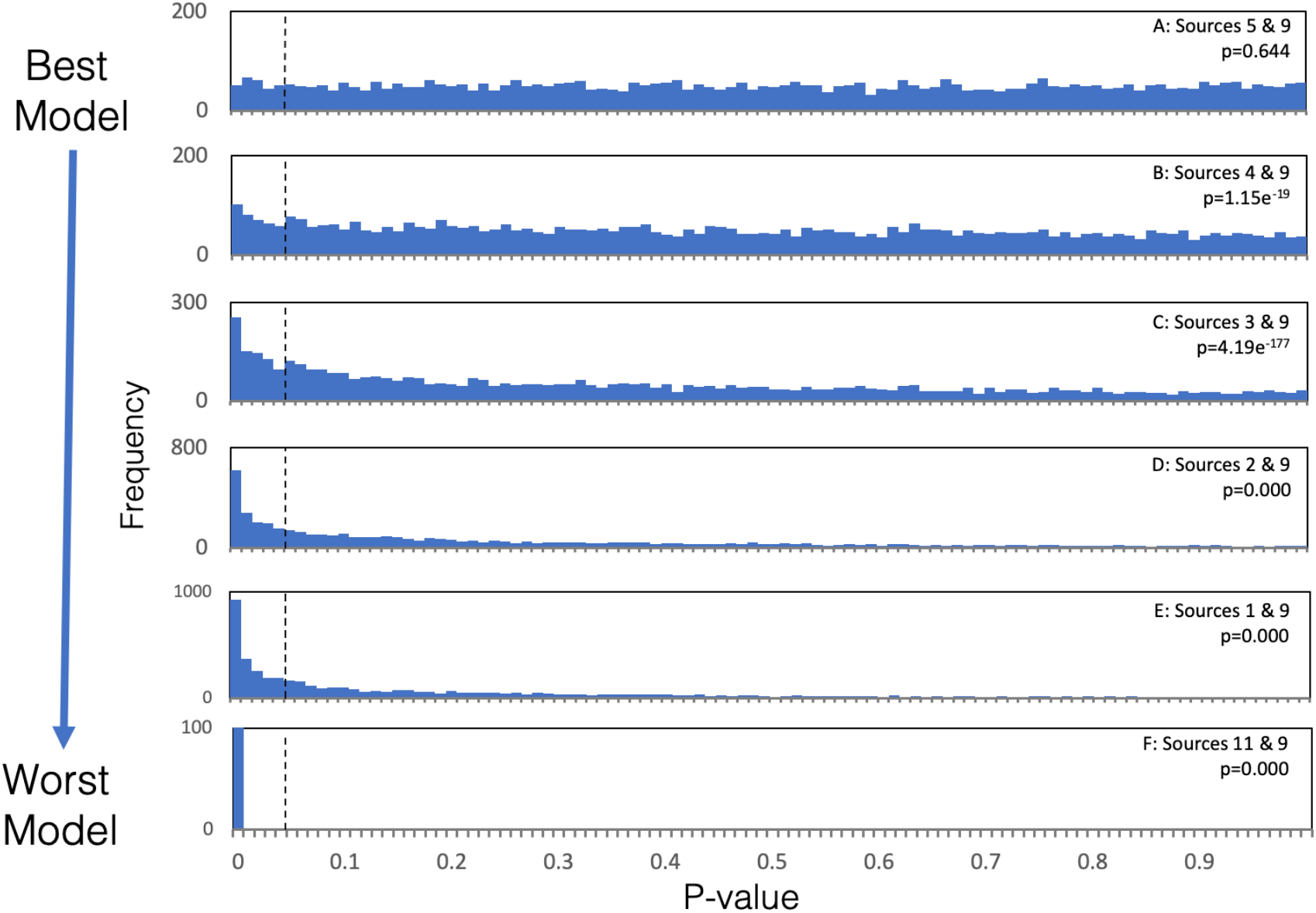
Distribution of p-values generated for various qpAdm models. The distribution of p-values generated by 5,000 replicates of qpAdm is shown for all models, except when sources 11 & 9 are used, in which case only 100 replicates were generated. Panel A shows the distribution of p-values produced by models using populations 5 & 9 as sources, which are the best possible sources of ancestry for population 14 out of the proposed models. Panels B-F show the distribution of p-values produced by models that use increasingly inappropriate source populations, relative to the chosen reference populations. Vertical black dotted lines indicate the p-value threshold of 0.05, above which qpAdm models are considered plausible. The results of a Kolmogorov-Smirnov test to determine whether the p-values are uniformly distributed are indicated.

As qpAdm is often used to distinguish between optimal and non-optimal models of admixture, we also seek to confirm that the distribution of p-values is not uniform when an incorrect model is considered. We therefore examine the distribution of p-values produced when non-optimal populations (i.e. populations 1-4 and 11) are used as sources instead of population 5. As populations 1-4 share more genetic drift with reference population 0 than the true source population (and similarly because population 11 shares less drift with population 0 than the true source population), we expect that the distribution of p-values produced by qpAdm should be biased towards zero when these populations are used as sources (with population 11 producing the strongest bias). We ran these non-optimal qpAdm models on the 5,000 replicate datasets described above and observe a deviation from a uniform distribution. In the case of populations 1-4, models that include source populations that share the most drift with population 0 yield p-value distributions that are most strongly biased towards zero (Figure 2B-F; Supplementary Table 1), and as expected, p-values associated with using population 11 as a source are even more strongly biased towards zero. In each case, using a Kolmogorov-Smirnov test, we reject the null hypothesis that the p-values are uniformly distributed.

Although the distributions of p-values deviate from a uniform distribution as expected, we also note that in the cases where populations 1-4 are used as potential source populations, a large proportion of these models are assigned p-values that would be considered plausible using 0.05 as a standard threshold. These results reflect the fact that populations 1-5 are all closely related (average pairwise F_ST_ between <0.001-0.005), therefore the inclusion of population 0 as the only reference population with the power to distinguish between these populations (as it is differentially related to them), may not be enough to reject models that use populations 1-4 as sources in all cases. In practice, if populations 1-5 were all proposed as potential sources and qpAdm assigned plausible p-values to multiple models, further analysis would be required to distinguish between these models. Further, we do note that when population 0 is excluded from the reference population set, all of the tested qpAdm models using populations 1-5 as a potential sources, produce approximately uniformly distributed p-values, as would be expected theoretically, as populations 1-5 are all symmetrically related to all other reference populations (Supplementary Figure 1; Supplementary Table 2).

While the overall distributions of p-values differ between optimal and non-optimal qpAdm models, we note that for individual replicates the most optimal model is not necessarily assigned the highest p-value. We find that the p-value associated with the best model (sources 5 & 9) produces the highest p-value in only 48% of cases (Supplementary Table 1). Therefore, in cases where multiple models are assigned plausible p-values (i.e. p ≥ 0.05), we caution that p-value ranking (i.e. selecting the model that is assigned the highest p-value) should not be used to identify the best model. Methods for distinguishing between multiple models will be discussed further in the section on comparing admixture models.

### Accuracy of Admixture Proportion Estimates

In addition to generating informative p-values, it is essential that qpAdm generates accurate admixture proportion estimates. This has also not been formally tested using simulated data. We therefore simulate genetic data according to the population tree shown in Figure 1, varying the proportion of admixture (α) occurring in the lineage ancestral to population 14 between 0.0-1.0 at intervals of 0.1 with 20 replicates per interval. We find that the estimated admixture proportions are extremely close to the actual simulated admixture proportions for all values of α (Figure 3A; Supplementary Table 3). In 99.3% of cases (220 total), the estimated α is within 3 standard errors of the simulated α, consistent with theoretical expectations, with an average standard error of 0.0092 [range: 0.008-0.011]. These results indicate that qpAdm accurately estimates admixture proportions, regardless of the level of admixture, and that the standard errors produced by qpAdm are well calibrated. However, we recognize that in practice, users of qpAdm have access to a much less complete dataset. Therefore, we modify the data in order to explore the performance of qpAdm when applied to data of lower coverage and quality.

**Figure 3.**
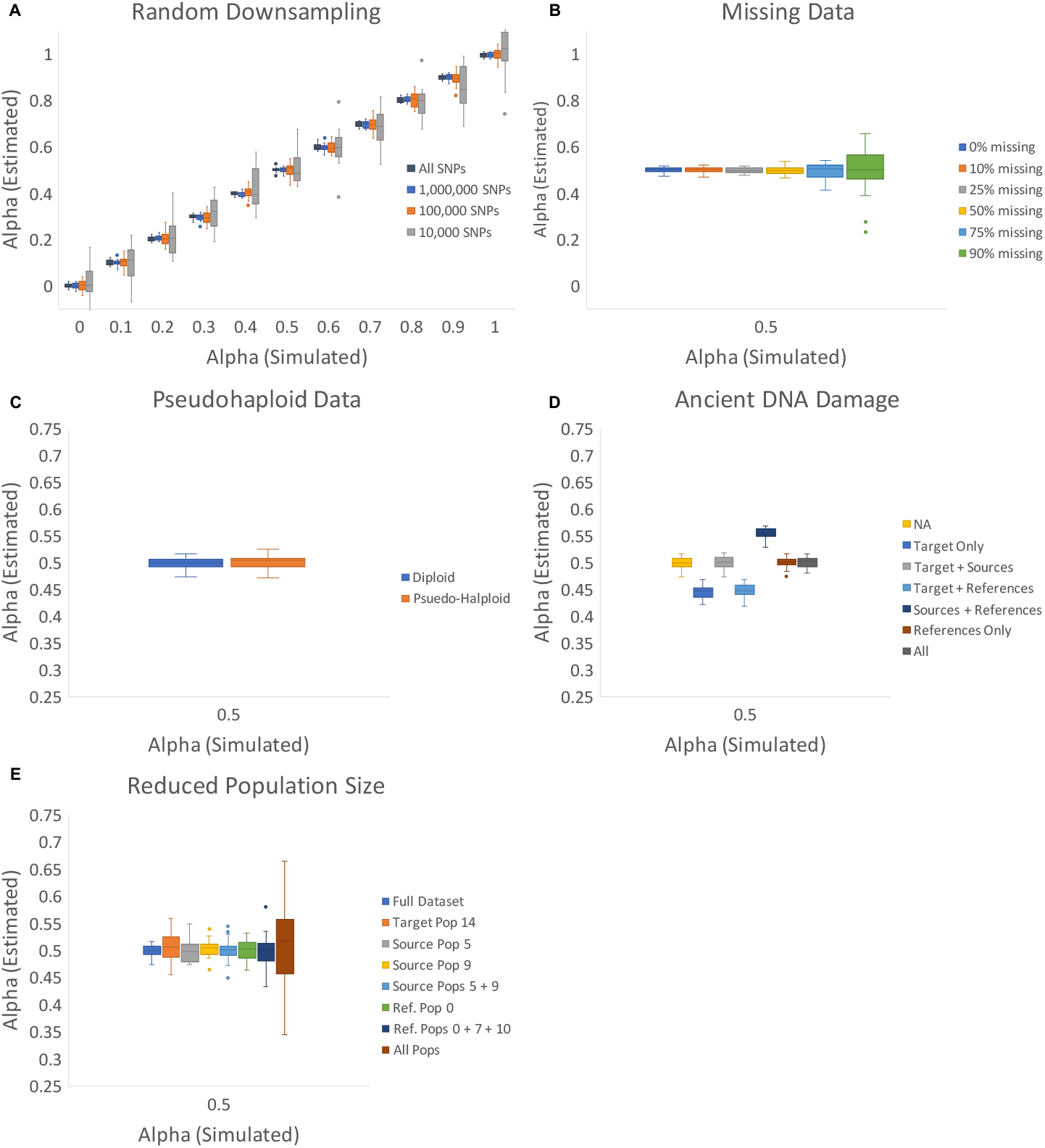
Accuracy of Admixture Proportion Estimates. Box and whisker plots showing the estimated values of admixture proportion (alpha) generated by qpAdm for varying simulated alphas. Only alpha 0.5 is shown for panels B-E, however all alphas 0-1 are reported in the corresponding Supplementary Tables. For each simulated alpha, 20 replicates of qpAdm are performed for each condition. [A] Estimates produced by qpAdm when run on the entire dataset and after randomly down-sampling to 1 million, 100 thousand, and 10 thousand SNPs. All subsequent analyses are performed on the 1 million SNP downsampled dataset [B] Estimates produced by qpAdm where some proportion (0%, 10%, 25%, 50%, 75% or 90%) of data is missing in each individual. [C] Estimates produced by qpAdm in both diploid and pseudo-haploid form. [D] Estimates produced by qpAdm where 5% ancient DNA damage is simulated in a subset of populations (14, 14+5+9, 14+0+7+10+12+13, 5+9+0+7+10+12+13, 0+7+10+12+13, and All populations). [E] Estimates produced by qpAdm, where only a single individual is sampled from varying populations (14, 5, 9, 5+9, 0, 0+7+10, and all populations).

Each simulation contains an average of ~30 million SNPs. In order to understand the performance of qpAdm with less data, we randomly down-sample the complete dataset to produce analysis datasets of 1 million, 100 thousand, and 10 thousand sites. In all cases, the average admixture proportion estimate generated is extremely close to the simulated α, although we do observe an increase in the amount of variance in the individual estimates as the amount of data analyzed decreases (Figure 3A; Supplementary Table 3). In order to increase computational efficiency and to better approximate typical analysis datasets, all subsequent analyses are performed on the data that has been randomly down-sampled to 1 million sites. We observe similar results when using non-random ascertainment schemes to select sites for analysis (Supplementary Table 4). The impact of non-random ascertainment schemes on qpAdm analyses are described in more detail in a later section.

We find that qpAdm is robust to missing data, where data from randomly selected sites in each individual is considered missing with rate 10%, 25%, 50%, 75% or 90% (Figure 3B; Supplementary Table 5). Additionally, we find that pseudohaploidy—a common feature of ancient DNA, where due to low sequencing coverage, a haploid genotype is determined by randomly selecting one allele at each diploid site and assigning that to be the genotype—has little impact on admixture estimates (Figure 3C; Supplementary Table 6).

Ancient DNA is also subject to deamination, resulting in C-to-T or G-to-A substitutions appearing in transition sites. In the 1.2 million SNP sites that are commonly targeted in ancient DNA analysis, approximately 77.6% of sites are transitions (Fu *et al.* 2015; Haak *et al.* 2015; Mathieson *et al.* 2015). We therefore randomly defined 77.6% of simulated sites to function as transitions. For each of these transition sites, in each individual, if the allele at that position is of the reference type, it was changed to the alternative type with 5% probability, mimicking the unidirectional change in allelic state caused by ancient DNA damage. We find that admixture proportion estimates produced by qpAdm are relatively robust to the presence of ancient DNA damage in cases where all populations exhibit an equal damage rate (Figure 3D; Supplementary Table 7). However, in cases where the target (population 14) and source (5 + 9) populations have a different rate of ancient damage the estimated admixture proportions are biased. This bias reflects attraction between populations on the left and right sides of the f_4_-statistics calculated by qpAdm and is not unexpected. The effects of differential rates of ancient DNA damage between populations on qpAdm analyses are explored further in a later section.

Another concern that is common among ancient DNA analyses is small sample size. We therefore explore the effect of reducing the sample size of various populations in the analysis from ten individuals down to a single individual. We find that admixture estimates are relatively robust to this reduced sample size regardless of whether the target (population 14), source (population 5, 9, or 5 + 9) or reference (population 0 or 0 + 7 + 10) set has only a single individual sampled (Figure 3E; Supplementary Table 8). Reducing the target sample size to a single individual appears to have the greatest effect for all cases where only the sample size of a single population was reduced. Further, we see that when only a single individual is sampled from every population, the admixture proportion estimates vary the most between replicates, however, the mean of these estimates fall close to the true α, suggesting that small sample size does not result in an upward or downward bias in the admixture proportion estimates produced by qpAdm.

While none of the factors considered here result in biased admixture proportion estimates (except for when ancient DNA damage is present non-uniformly across populations), we caution that the increase in variance associated with each of these forms of reduced data quality is likely to be additive, so models relying on data with high rates of missingness, damage and very small sample sizes should be interpreted with caution.

### Comparing Admixture Models

One of the major applications of qpAdm is to identify an optimal admixture model, out of a variety of proposed possible models, many of which may be deemed plausible by qpAdm. However, no formal recommendations have been made about what strategy to use when comparing models. We therefore explore two commonly employed approaches for comparing admixture models in order to make recommendations for best practices in qpAdm usage.

One of the most typical implementations of qpAdm involves the selection of a set of differentially related populations to serve as the base set of reference populations. This base set of reference populations is often chosen to represent key positions in the known population history (i.e. the ‘O9’ reference set defined in Lazaridis *et al.* (2016)). A non-overlapping set of source populations is then defined, and qpAdm models involving different combinations of source populations and the base set of reference populations are tested. Using this method, multiple models may meet the criteria to be considered plausible, and the most optimal model is identified by adding additional reference populations to the base set of references, which are selected for their differential relatedness to one or more of the source populations in the set of potentially plausible qpAdm models.

While this strategy is relatively straightforward and widely implemented (e.g. Lazaridis *et al.* 2016; Harney *et al.* 2018), it has several drawbacks. In particular, because a population cannot simultaneously serve as a source and reference population, this strategy either requires that populations that are placed in the base set of reference populations are not considered as potential source populations (meaning it is possible that the best source population would be entirely missed if it were selected to serve in the reference population set) or that potential source populations be selectively removed from the reference population set so that they can be used as source populations for some models. This strategy results in the creation of some models that are not equivalent, and therefore are difficult to compare.

An alternative to the “base” reference set strategy that has been implemented in order to avoid these problems is to create a set of “rotating” models in which a single set of populations is selected for analysis (e.g. Skoglund *et al.* 2017; Harney *et al.* 2019). From this single set of populations, a defined number of source populations are selected, and all other populations then serve in the reference population set for the model. Under this “rotating” scheme, populations are systematically moved from the set of reference populations to the set of sources. Thus, all population models are generated using a common set of principles and are therefore more easily directly compared. In order to compare the performance of these two strategies (“base” versus “rotating”), we again focus on the population history of population 14 (Figure 1).

For the “base” reference approach, we continue to use the base set of reference populations as previously defined (populations 0, 7, 10, 12, and 13), all other populations are considered to be potential source populations. We used qpAdm to test all possible combinations of two source populations. We ran each of these qpAdm models on the data generated using the standard population history with α=0.50, with 20 replicates. Among these 20 replicates, qpAdm identified the optimal model, in which populations 5 and 9 serve as source populations, as plausible in 19 cases (Figure 4; Supplementary Table 9). However, there are also a large number of other population models that are consistently deemed plausible; for example when population 8 is used as a source (in conjunction with population 5) instead of population 9, 90% of the models are deemed plausible. The high rate of acceptance of this model is fully consistent with expectations, because while population 9 is more closely related to the true source population, populations 8 and 9 are symmetrically related to all of the reference populations included in the model, and therefore are indistinguishable using this approach (unless data from a population that differentially related to these two populations could be added to the model). Models that include populations 1-4 (in combination with populations 8 or 9) were also frequently identified as plausible. These results suggest that the inclusion of population 0 as a reference does not provide enough information to differentiate between these potential source populations and the true optimal source (population 5). Therefore, the next step in a qpAdm analysis that utilizes the base model approach would be to add additional reference populations that are differentially related to populations 1-5 in order to help differentiate between the remaining possible models.

**Figure 4.**
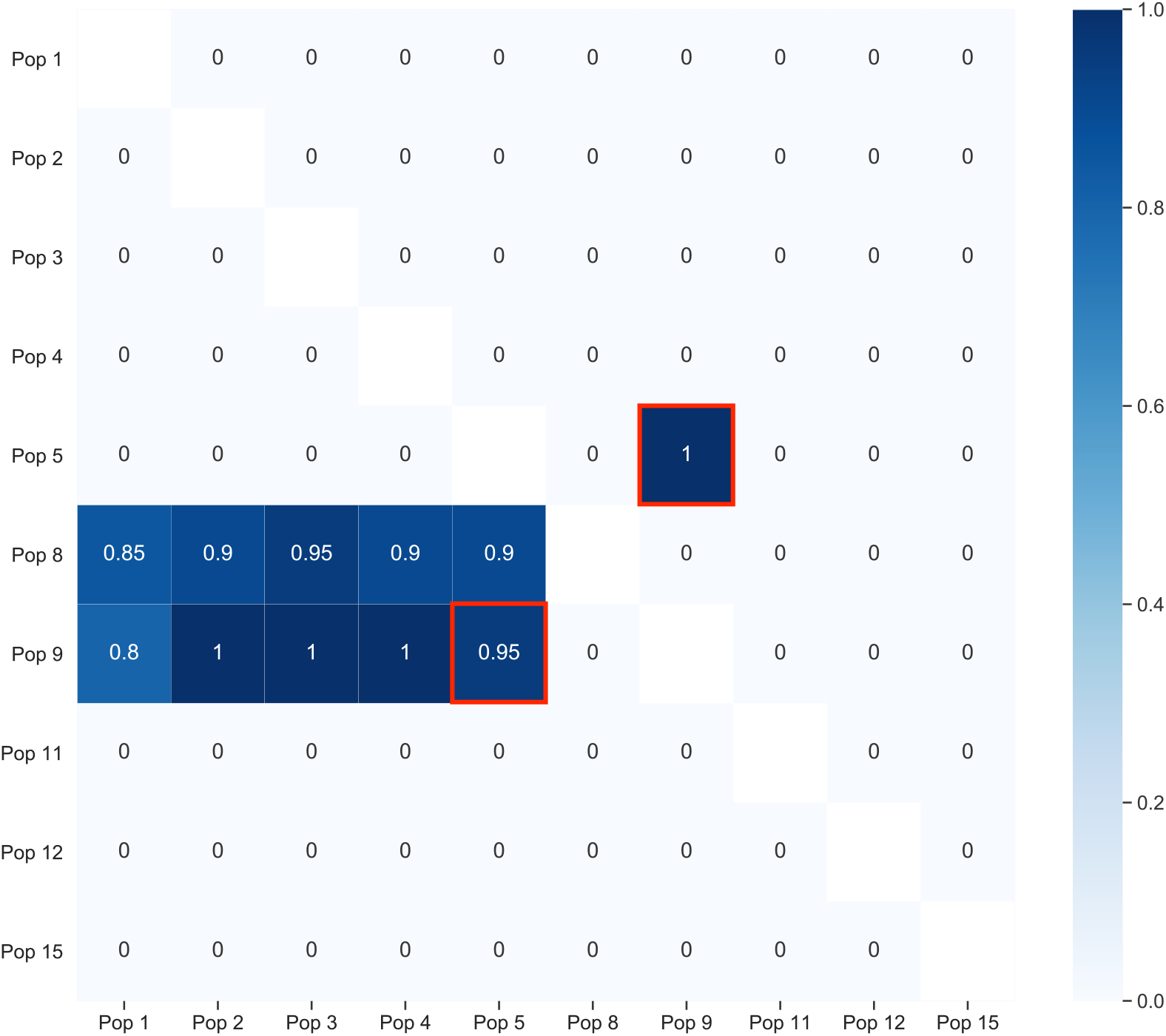
Comparing qpAdm models using various approaches. A heatmap showing the proportion replicates in which the 2-way admixture model generated using each combination of possible source populations is deemed plausible by qpAdm (i.e. yielded a p-value > 0.05 and admixture proportion estimates between 0-1). The lower triangle shows results generated using the base set of reference populations (0, 7, 10, 12, and 13), while the upper triangle shows results generated the rotating model approach. The proportion of replicates deemed plausible is indicated by the color (darker shades indicate a higher proportion) and is written inside each square of the heatmap. The optimal admixture model(s) for each of the approaches are highlighted in red.

In contrast, we find that under a “rotating” model, where all populations (except for population 6 because it is phylogenetically a clade with source 5) were selected to serve as either a source or a reference population, all models that included populations 5 and 9 as sources were identified as plausible. In contrast all other population models were rejected (Figure 4; Supplementary Table 10). The inclusion of the optimal source populations (5 & 9) as references in all other models enables qpAdm to differentiate between models that would otherwise be indistinguishable (such as differentiating between populations 8 and 9 and between populations 1-5). Further, in cases where optimal source populations are not available (i.e. if both populations 5 and 6 are excluded from the model), qpAdm still identifies closely related models as plausible (such as those involving admixture between population 9 and populations 0-4), suggesting that this rotating approach is not overly stringent in cases where optimal data are not available (Supplementary Table 11). Due to the relative simplicity of the rotating model approach and the increased ability to identify the optimal admixture model when using it, we recommend utilizing a rotating strategy when possible.

#### Ascertainment bias and “rotating” model selection

In order to understand the impact of ascertainment bias on model selection, we repeated this analysis on data that was ascertained from the full dataset using several non-random ascertainment strategies, including ascertaining on sites that were found to be heterozygous in a single individual from [1] the target (population 14), [2] a source (population 5), and [3] two populations that are uninvolved in the admixture event (population 10 and 13). The individual used for data ascertainment was excluded from subsequent analyses. In all cases, using the rotating approach previously described, only models that use populations 5 and 9 as sources are deemed plausible (Figure 5; Supplementary Table 12), suggesting that ascertainment bias is unlikely to cause users to identify inappropriate models as plausible. Further, the optimal model was identified as plausible in at least 90% of replicates using all ascertainment strategies, suggesting that qpAdm is robust to ascertainment bias. These results are consistent with previous findings that f_4_-statistics, which are used for all qpAdm calculations, are robust to biased ascertainment processes (Patterson *et al.* 2012).

**Figure 5.**
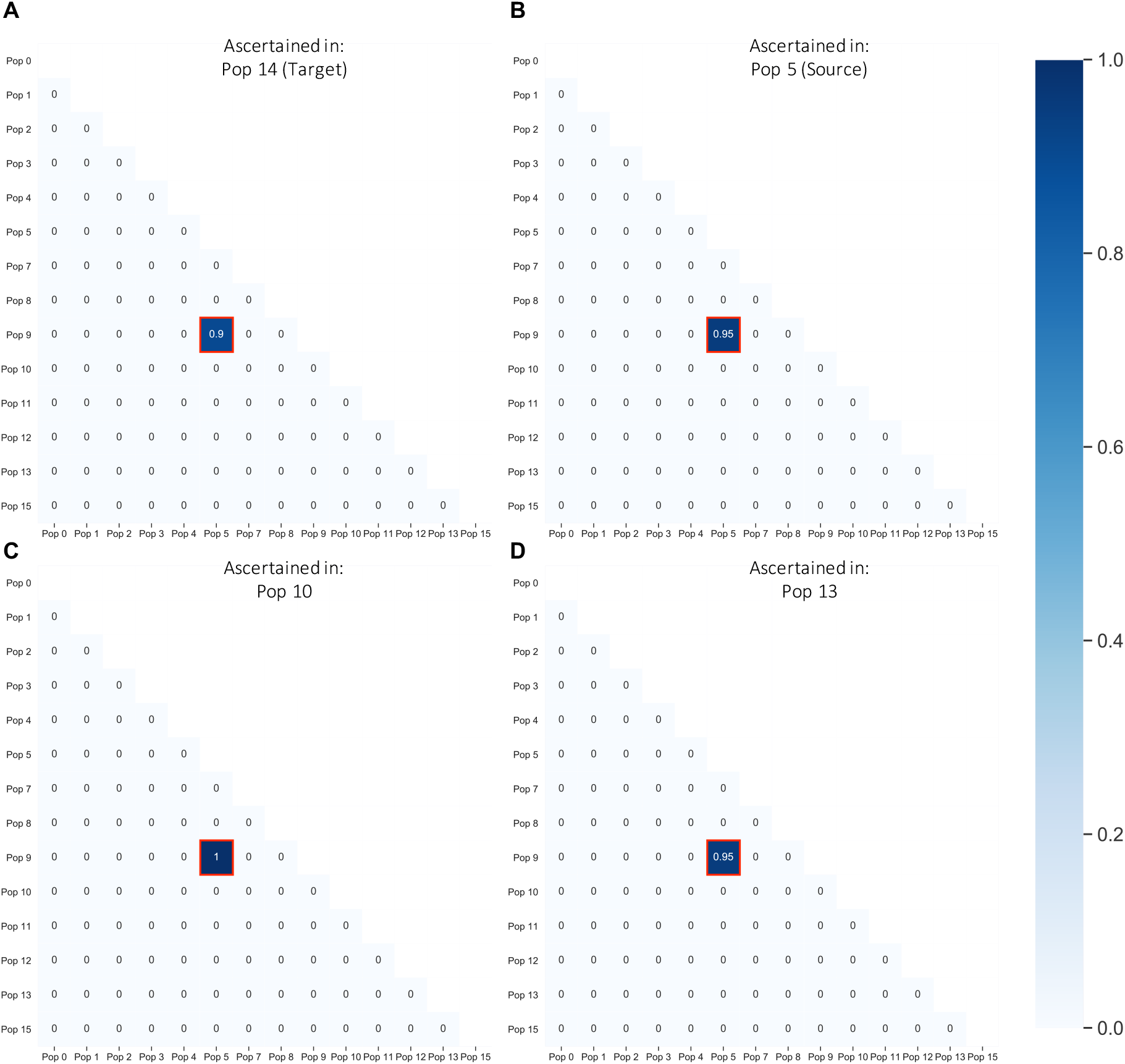
Effect of ascertainment bias on qpAdm model selection. Heatmaps showing the proportion of replicates in which the 2-way admixture model generated using each combination of possible source populations is deemed plausible by qpAdm (i.e. yielded a p-value > 0.05 and admixture proportion estimates between 0-1) on SNP data that is ascertained from a heterozygous individual in a single population, [A] population 14 (target), [B] population 5 (source), [C] population 10 and [D] population 13. The proportion of replicates deemed plausible is indicated by the color (darker shades indicate a higher proportion) and is written inside each square of the heatmap. The optimal admixture model for each of the approaches are highlighted in red.

#### Missing data and the “allsnps” option of qpAdm

We also explored the effect of qpAdm’s “allsnps” option when working with samples with a large amount of missing data. If the default “allsnps: NO” option is selected, qpAdm only analyzes sites that are shared between all target, source, and reference populations that are included in the model. In contrast, if “allsnps: YES” is selected, every individual f_4_-statistic is calculated using the intersection of SNPs that have available data for the four populations that are involved in that particular calculation, therefore every f_4_-statistic is calculated using a unique set of sites. The “allsnps: YES” parameter is commonly used in cases where one or more populations in the analysis dataset has a high rate of missing data, in order to increase the number of sites analyzed. However, this causes the underlying calculations performed by qpAdm to deviate from those on which the theory is based, and the effect of this change in calculations on admixture proportions estimated by qpAdm and on optimal model identification is not well studied.

We explore the effects of this parameter, using simulated data with admixture proportion α=0.50 and rates of missing data equal to either 25%, 80%, 85% or 90% for all individuals across 1 million SNP sites. We implemented the rotating model for both the “allsnps: YES” and “allsnps: NO” options (all previous analyses used the “allsnps: YES” option). Comparing all possible models using the rotating approach, we find that the results produced when using the “allsnps: YES” and “allsnps: NO” options are similar when the rate of missing data is low (i.e. 25%) (Figure 6A; Supplementary Table 13). The optimal model (with sources 5 and 9) was identified as plausible in 95% of cases and no other models were deemed plausible for both options. Further, the admixture proportion estimates produced in both cases are relatively similar, with average standard errors of 0.006 in both cases. The similar performance of the “allsnps: YES” and “allsnps:NO” options in this case is likely due to the relatively large sample size (10 individuals per population) used in the analysis. With 25% missing data, the expected number of SNPs to be included the analysis when the “allsnps: YES” option is selected is 1 million. This number is only slightly reduced, to 999,985.7, when the “allsnps: NO” option is selected.

**Figure 6.**
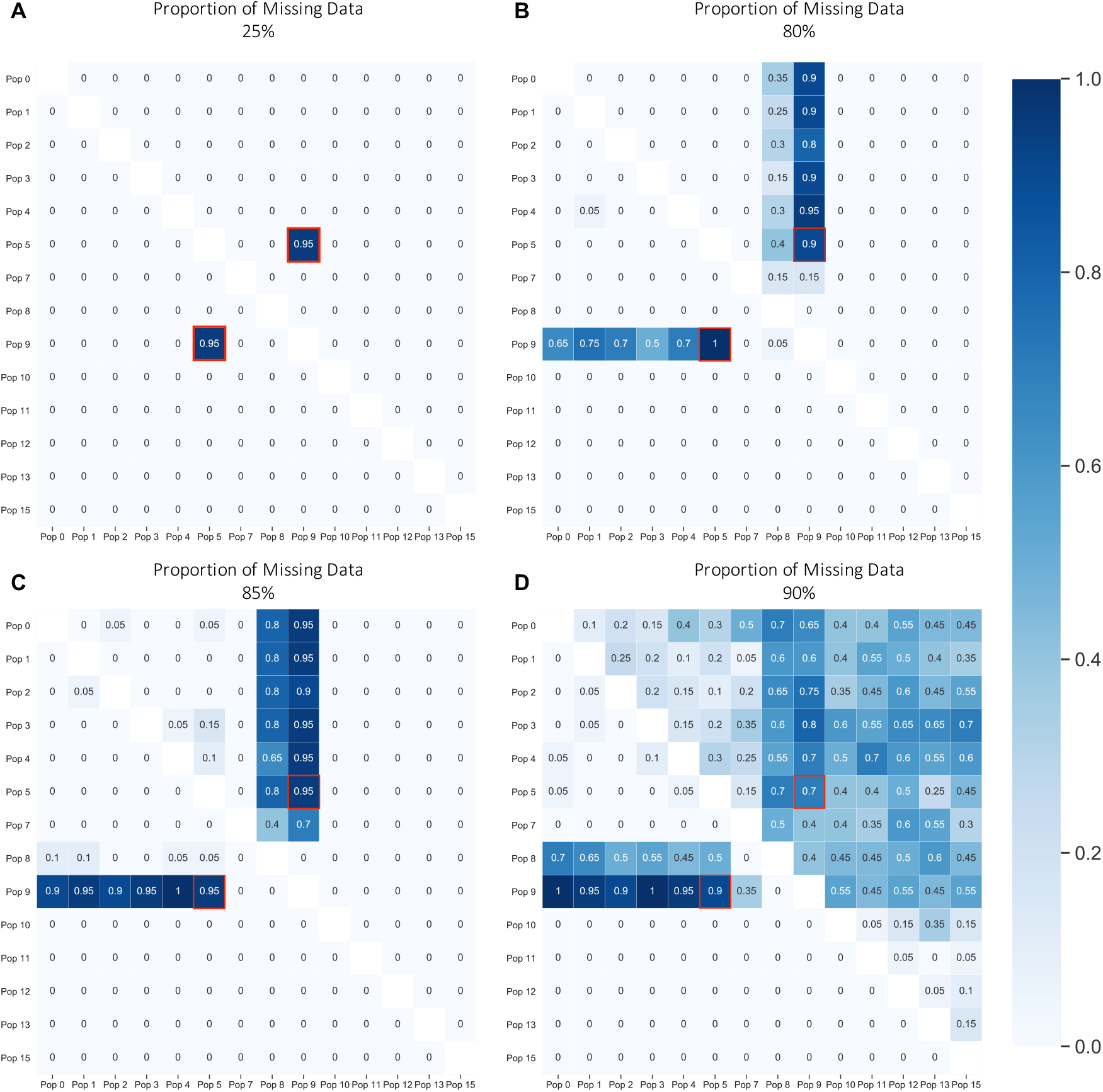
Effect of the allsnps parameter on qpAdm model selection. Heatmaps showing the proportion of replicates in which the 2-way admixture model generated using each combination of possible source populations is deemed plausible by qpAdm (i.e. yielded a p-value > 0.05 and admixture proportion estimates between 0-1) on SNP data using the “allsnps: yes” (lower left triangle) and “allsnps: no” parameters (upper right triangle), on data with [A] 25% [B] 80% [C] 85% or [D] 90% missing data. The proportion of replicates deemed plausible is indicated by the color (darker shades indicate a higher proportion) and is written inside each square of the heatmap. The optimal admixture model for each of the approaches are highlighted in red.

In contrast, when the rate of missing data is elevated (i.e. 80%, 85% or 90%), a difference in performance between the “allsnps: YES” and “allsnps: NO” options was observed. In each case, when the rate of missing data increased, the number of non-optimal models that were identified as plausible also increased (Figure 6B-D). These changes were more dramatic when the “allsnps: NO” parameter was used, further we observe a greater increase in the standard errors associated with admixture proportion estimates produced when using the “allsnps: NO” option, with average standard errors equal to 0.025, 0.066, and 9.994 when analyzing data with 80%, 85%, and 90% missing data, respectively. In contrast, while the standard errors produced using the “allsnps: YES” option also increased, the increase was lower in magnitude in all cases, with standard errors of 0.015, 0.020, and 0.035 observed, respectively. This difference in performance is likely the result of the number of SNPs available for analysis when using each option. When using the “allsnps: YES” parameter, the expected number of SNPs used in analysis of data with 80%, 85%, and 90% missing data rates remains 1 million. However, when using the “allsnps: NO” parameter, the expected number of SNPs used in analysis with each rate of missing data is only 181,987.5, 37,303.7, and 1,610.4 SNPs, respectively. These results suggest that the increased data provided by using the “allsnps: YES” option improves the ability of qpAdm to distinguish between models, without creating biases in cases where missing data is distributed randomly throughout the genome of all individuals.

#### The effects of ancient DNA damage on model selection

In an earlier section, we show that admixture proportion estimates produced by qpAdm can be biased when produced using populations with differential rates of ancient DNA damage. We therefore explored the effects of damage on model comparison, using the rotating model approach. Across all cases, only models involving the optimal sources (populations 5 and 9) are deemed plausible, suggesting that ancient DNA damage, even when unevenly distributed, is unlikely to cause a user to identify a non-optimal model as plausible (Figure 7; Supplementary Table 14). Further, when damage rates are consistent between the target and optimal source populations, the optimal model is identified as plausible in at least 95% of cases. However, when the target and source populations have differential rates of damage, this optimal model is almost always deemed implausible. We do note that the ancient DNA damage simulated in this analysis (5% ancient DNA damage rate at all “transition” sites) is relatively high, as most ancient DNA damage occurs at the terminal ends of DNA molecules. Therefore, these results likely represent an extreme case. However, these results highlight the importance of considering the effect of ancient DNA damage in ancient DNA analyses. In particular, we caution against designs where both ancient and present-day populations are included in a single qpAdm model.

**Figure 7.**
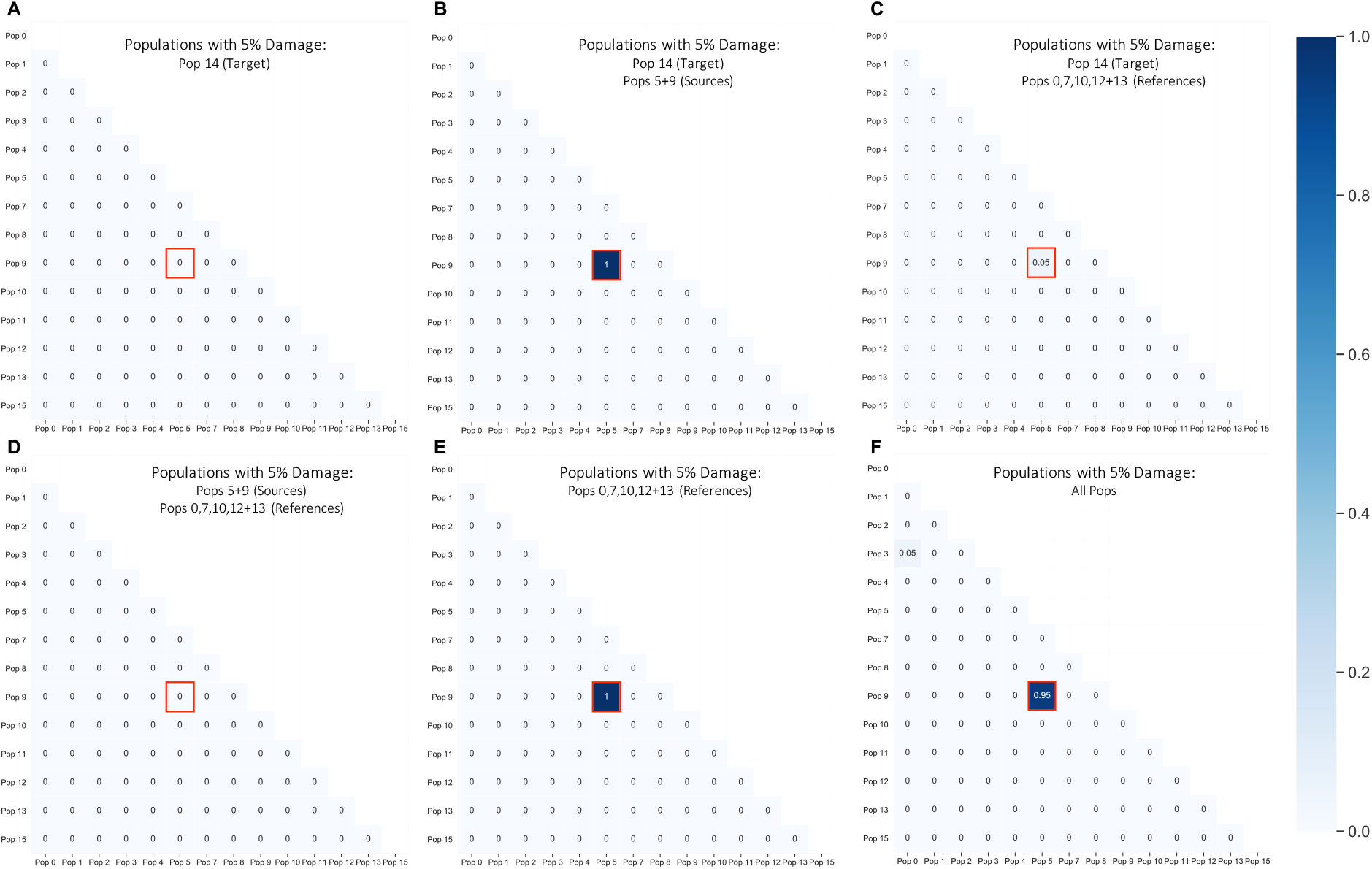
Effect of ancient DNA damage on model selection. Heatmaps showing the proportion of replicates in which the 2-way admixture model generated using each combination of possible source populations is deemed plausible by qpAdm (i.e. yielded a p-value > 0.05 and admixture proportion estimates between 0-1) on SNP data. In each case [A-F] a given population or set of populations (14, 14+5+9, 14+0+7+10+12+13, 5+9+0+7+10+12+13, 0+7+10+12+13 and all populations) contain ancient DNA damage at 5% of “transition” sites. The proportion of replicates deemed plausible is indicated by the color (darker shades indicate a higher proportion) and is written inside each square of the heatmap. The optimal admixture model for each of the approaches is highlighted in red.

#### The effects of sample size on model selection

We also considered the impact of limited sample size when comparing models, using a rotating model approach. Using the same data shown in Figure 3E, where the sample size of the specified population(s) was reduced to 1 (Figure 8; Supplementary Table 15). In cases where the population(s) with reduced sample size were not involved in the admixture event of interest the effect of sample size reduction is minimal. Similarly, the results do not appear to be significantly affected when population 9 (one of the optimal source populations) experiences reduced sample size, suggesting that when the optimal source population is relatively differentiated from all other populations considered, reduced sample size has little effect. However, when source population 5 only contained a single sampled individual, models using closely related populations as sources were also deemed plausible. Similarly, when the target population (14) contained only a single sampled individual, the proportion of non-optimal models that were identified as plausible by qpAdm increased. These results suggest that when the sample size is lower, particularly for target or source populations, qpAdm has less power to reject non-optimal models. This is likely to become an even greater issue in cases where populations included in qpAdm models contain only a single individual with large amounts of missing data.

**Figure 8.**
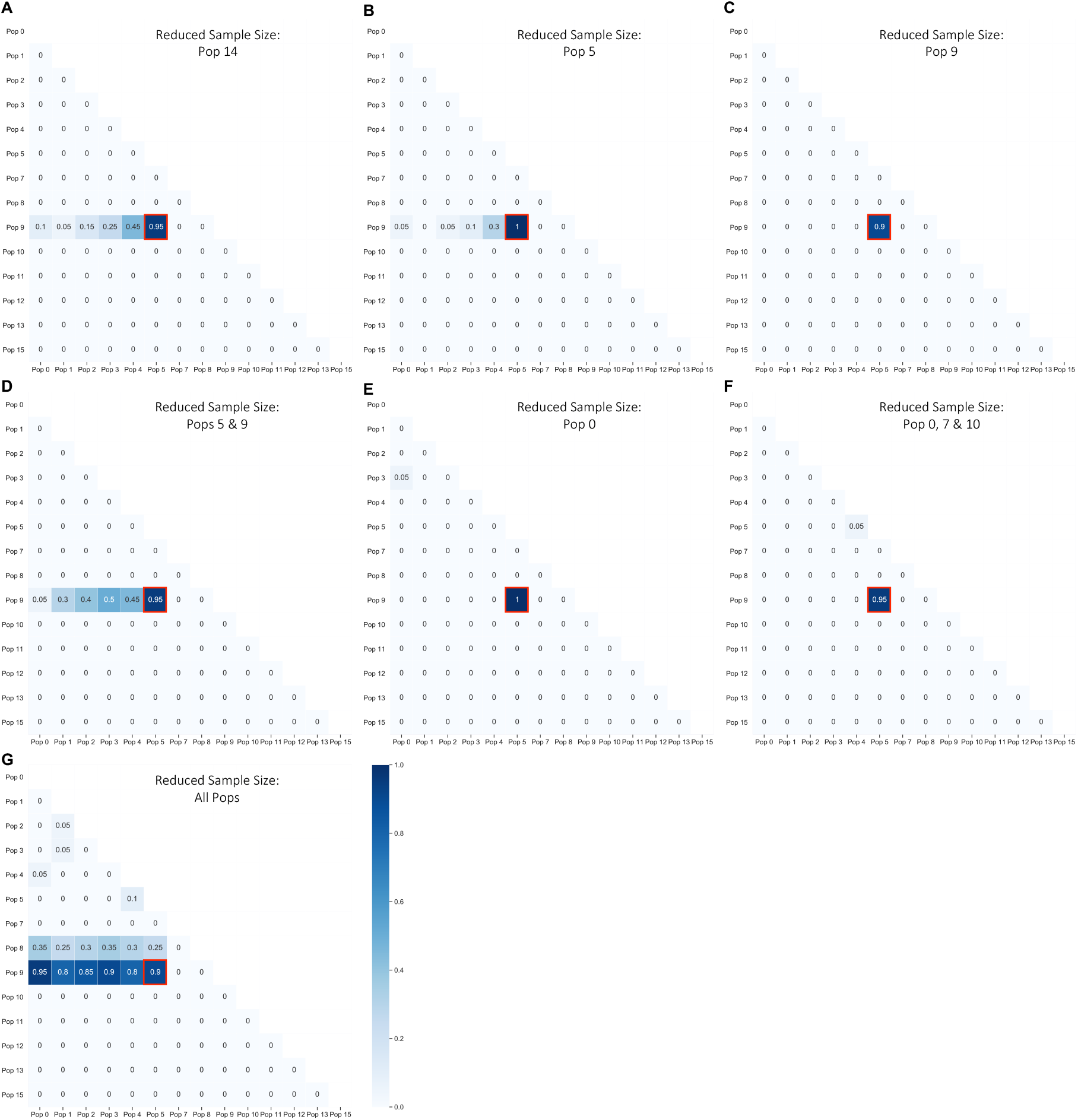
Effect of reduced sample size on model selection. Heatmaps showing the proportion of replicates in which the 2-way admixture model generated using each combination of possible source populations is deemed plausible by qpAdm (i.e. yielded a p-value > 0.05 and admixture proportion estimates between 0-1) on SNP data. In each case [A-G] a given population or set of populations (14, 5, 9, 5+9, 0, 0+7+10, and all populations) contain only a single sampled individual. The proportion of replicates deemed plausible is indicated by the color (darker shades indicate a higher proportion) and is written inside each square of the heatmap. The optimal admixture model for each of the approaches is highlighted in red.

#### Modeling unadmixed populations using qpAdm

Finally, while we know that the population history of population 14 involves admixture, the number of ancestral sources that contributed ancestry to a real target population is typically unknown. Therefore, we explored the behavior of qpAdm when modeling the population history of unadmixed and admixed populations (populations 6 and 14, respectively) under various scenarios. First, we explored models in which only a single source population contributed ancestry to the target population, using the same rotating model as described previously, but only selecting a single source population for each model. In the case of the unadmixed population 6, we find that in 95% of cases, it can be modeled as forming a genetic clade with population 5, consistent with theoretical expectations (Supplementary Table 16). In contrast, population 14 is never found to form a genetic clade with any of the tested source populations (Supplementary Table 17), again consistent with expectations. However, when population 6 is modeled as the product of admixture between 2 source populations, we find that it is frequently modeled as the product of a two-way admixture between population 5 and any other source population, where population 5 is estimated to contribute the vast majority of ancestry to population 6 (Supplementary Table 18)—in cases where these models are rejected, it is typically because population 5 is modeled as contributing greater than 100% of the ancestry to population 6, rather than due to a low p-value. We therefore stress the importance of testing all possible models with the lowest rank (i.e. number of source populations) using qpAdm (or the related qpWave) before proceeding to test models with higher rank.

### Challenging Scenarios

While we find that qpAdm behaves as expected under standard conditions, we are also interested in identifying scenarios under which qpAdm might behave in unanticipated and undesirable ways. We therefore explore the performance of qpAdm under two challenging scenarios: when the number of reference populations is very large and when the relatedness of populations is not tree-like but rather reflects ongoing genetic exchange.

#### Number of reference populations

We were interested in the effect of assigning an extremely large number of populations to the reference population set. While a commonly employed method for distinguishing between optimal and non-optimal admixture models and reducing the standard errors associated with a admixture proportion estimates is to increase the number of reference populations included in qpAdm models (e.g. Lazaridis *et al.* 2016; Harney *et al.* 2018), the effect of including too many reference populations in a model is unknown. As qpAdm generates f_4_-statistics involving combinations of reference populations, the larger the number of reference populations is, the more poorly estimated the covariance matrix of these f_4_-statistics is predicted to be. Therefore, existing guidelines for qpAdm usage recommend against assigning too many populations to the reference set, as the computed p-values are thought to be unreliable. However, how many reference populations is “too many” and what the effect of exceeding this number would be on the calculated p-values is unknown.

We therefore simulated a dataset with a large number of populations by adding two additional population branching events, occurring 50 generations apart, to all locations on the standard population tree that are marked with a star in Figure 9A, resulting in a total of 118 total populations in the simulated dataset (see Supplementary File 2 for exact simulation parameters). After down-sampling the simulated data to 1 million sites, we then ran qpAdm, with population 14 as the target, and populations 5 and 9 as sources. Populations 0, 7, 10, 12 and 13 were again assigned to serve as reference populations. All other populations (excluding population 6) were added, one at a time in random order to the reference population set, resulting in qpAdm models with between 5 and 114 reference populations. As each new reference population was added to the model, we re-ran qpAdm and recorded the p-value.

**Figure 9.**
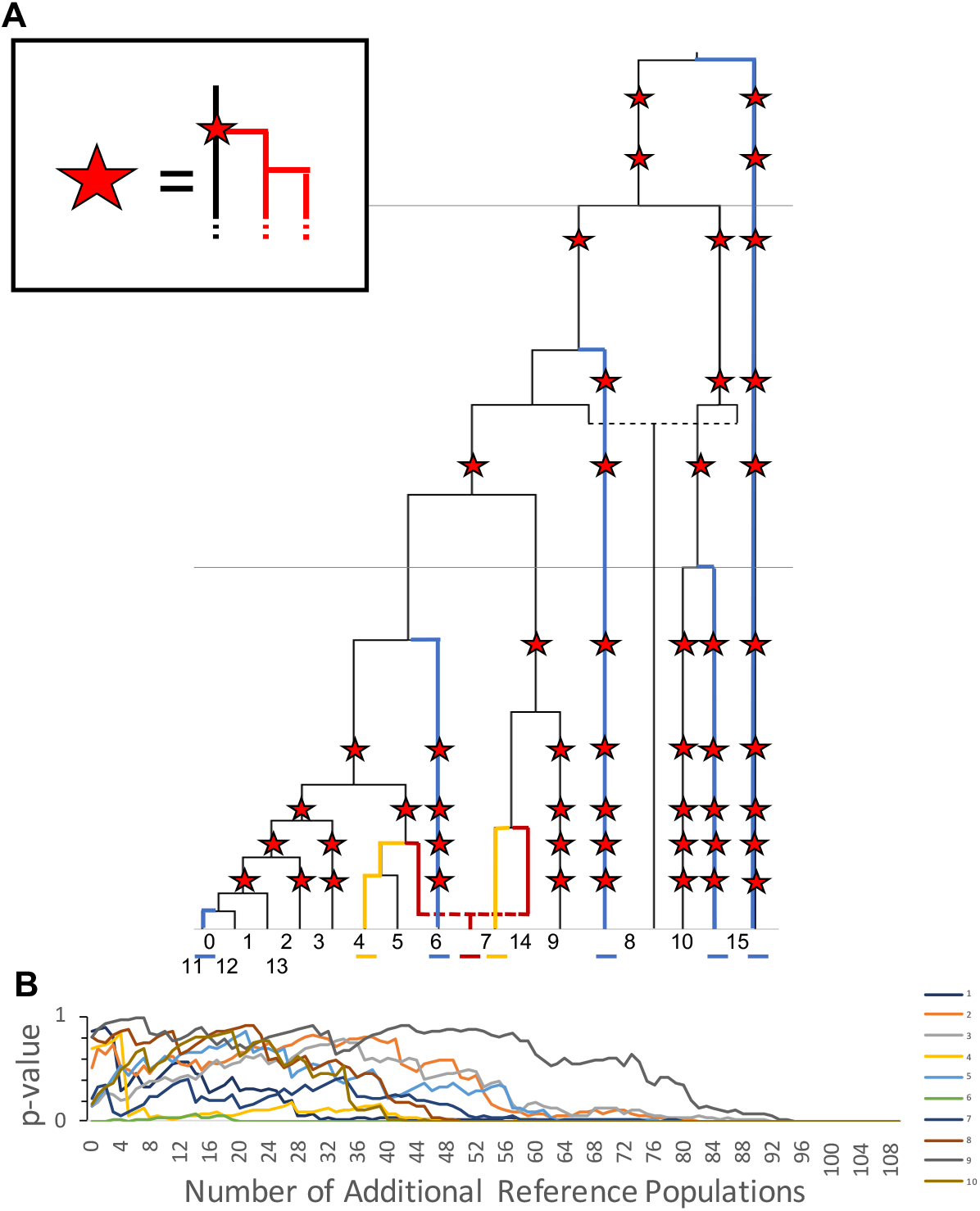
Inclusion of a large number of reference populations. (A) Population history of simulated data with additional populations added to tree. In all positions in the population history marked by a star, a population branching event occurs, forming an additional population. This new lineage undergoes an additional population branching event 50 generations later, resulting in two new populations created at each location marked with a star. Colors indicate the populations used in the base model, with the target in red, sources in yellow, and initial references shown in blue. (B) The change in p-values assigned to each model by qpAdm as additional reference populations are randomly added to the model. Each line tracks the p-values assigned to a single replicate, as the number of additional reference populations added to the base set of reference populations increases from 0 to 100.

Figure 9B shows the change in estimated p-value as reference populations are added to the model for 10 separate replicates (Supplementary Table 19). While the p-values calculated for each replicate using the original set of 5 reference populations appear to fall randomly between 0-1 (consistent with the uniform distribution of p-values observed in earlier analyses), we find that in all cases, as the number of reference populations increases the p-values eventually fall below the threshold of 0.05, resulting in all of the models with the maximum number of reference populations to be rejected. These results indicate that the inclusion of too many reference populations is likely to result in the rejection of qpAdm models, even in cases where the optimal source populations have been specified.

The maximum number of reference populations that can be included in a qpAdm model before this effect is observed is likely to depend on the specific population history and the total amount of data included in the analysis. In these simulations, we find that qpAdm begins to reject models that would otherwise be deemed plausible when as few as 30 additional populations are added to the outgroup set. These results support previous warnings against including too many reference populations in qpAdm models.

#### Continuous gene flow

An underlying assumption of qpAdm is that population admixture occurs in a single pulse over a small interval of time, during which the proportion of ancestry coming from each of the ancestral source populations can be estimated. However, real population histories often involve continuous gene flow that occurs over a prolonged period of time. In this case, although the resulting population may have received ancestry from multiple sources, estimates of admixture proportions from these sources may not be meaningful.

We therefore consider data simulated using a stepping-stone model of migration, in which neighboring populations exchange migrants each generation with rate, *m* (Kimura and Weiss 1964). We simulated a population history based on this migration model (Figure 10A), where 6 populations (each with an effective population size of 10,000) split from a common ancestral population 1000 generations previously, after which point migration occurred between neighboring populations. The model also includes three additional populations that are symmetrically related to these 6 populations, with all 9 lineages splitting from a common ancestral population 2000 generations in the past (see Supplementary File 3 for exact simulation parameters).

**Figure 10.**
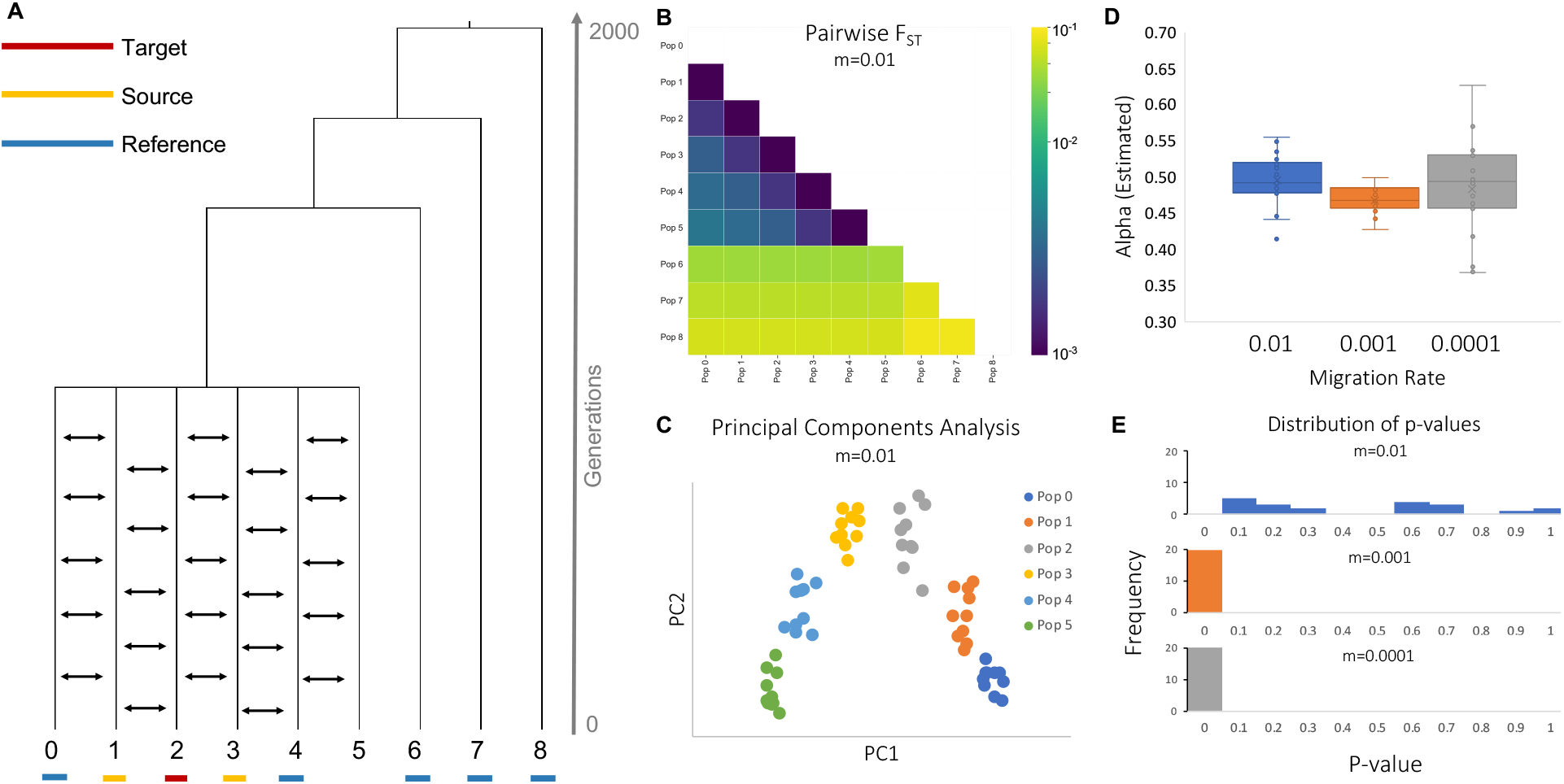
Continuous migration models. (A) Population history involving continuous migration. The target, source, and reference populations underlined in red, yellow, and blue, respectively. (B) A heatmap showing average pairwise FST between each population for 20 replicates (C) A PCA plot showing the relationship between all populations, calculated using a single replicate (D) Admixture proportions assigned by qpAdm for a model with population 2 as the target, and populations 1 and 3 as sources at varying migration rates. (E) Histograms showing the frequency of p-values produced by this qpAdm model at varying migration rates.

While under this model, populations 1 and 3 have each contributed ancestry to population 2, it would be inaccurate to say that population 2 is the product of admixture between these two populations. The duration of exchange of ancestry is much longer than what is supposed in qpAdm. In addition, population 2 was formed in the same population-splitting event that formed populations 1 and 3, not as the result of admixture between distinct populations 1 and 3. Finally, by symmetry population 2 is just as much the source of populations 1 and 3 as either of these is the source of population 2.

Preliminary analyses of the relationships between these 9 populations using pairwise F_ST_ (Patterson *et al.* 2006) would indicate that population 2 is closely related to both populations 1 and 3 (Figure 10B; Supplementary Table 20). Further, if populations 0-5 are plotted using PCA (Figure 10C; Supplementary Table 21)(Patterson *et al.* 2006), population 2 appears to fall on a genetic cline between these two populations. These results could be interpreted as suggestions that population 2 is the product of admixture between populations 1 and 3. While it might be possible using other *f*-statistics to determine that the relationship between these populations is not well described by a pulse admixture event (Lipson 2020), there is nothing to prevent a naïve user from attempting to model this relationship as the product of admixture using qpAdm. We therefore explore the effects of attempting to model the ancestry of population 2 (the target population) as the product of admixture between populations 1 and 3 (the source populations), with populations 0, 4, 6, 7 and 8 classified as reference populations.

We first consider the case of a very high migration rate (*m*=0.01; equivalent to 100 migrants moving from one population to the neighboring population per generation). Out of 20 replicates, qpAdm identifies the proposed model as plausible in 90% of cases, suggesting that qpAdm cannot always distinguish between population histories that involve continuous migration and those involving pulses of admixture. Further, qpAdm assigns admixture proportions of approximately 50% to each source population, which is sensible because each population does contribute roughly equal amounts of ancestry to the target population (Figure 10D-E; Supplementary Table 22). When we consider lower migration rates (m=0.001 and m=0.0001) we observe similar admixture proportion estimates, but all of the p-values fall well below the 0.05 threshold, suggesting that with lower rates of migration, qpAdm will reject admixture as a plausible model when the actual history involves continuous migration.

These results suggest that users should be sure to consider alternative demographic models to pulse admixture, even in cases when qpAdm produces admixture proportion estimates and p-values that appear plausible. This scenario likely represents just one of many cases in which qpAdm identifies plausible admixture models for populations that were not formed via admixture, therefore, we caution that users should use additional tools, in conjunction with or prior to qpAdm analysis, to determine whether admixture is a likely demographic scenario.

## DISCUSSION

We find that qpAdm can accurately identify plausible admixture models and estimate admixture proportions when applied to simulated data, matching previous theoretical expectations (Haak *et al.* 2015). When an appropriate admixture model is suggested, qpAdm calculates p-values that follow a uniform distribution, suggesting that a cut off value of 0.05 will result in the acceptance of a correct model in 95% of cases. Additionally, qpAdm estimates admixture proportions with high accuracy, even when calculated on datasets with a limited number of SNPs, high rates of missingness or damage (when occurring at similar rates in all populations), or when analyses are performed on pseudo-haploid data or on data that is subject to strong ascertainment bias. Additionally, while the use of populations with small sample sizes does increase the variance in admixture proportion estimates, admixture proportion estimates appear unbiased.

Further, we tested two commonly used strategies for identifying the best admixture model using qpAdm—base and rotating—and find that both strategies can distinguish between plausible and implausible models. However, the rotating strategy is better able to distinguish between plausible and implausible models, particularly when the potential source populations are closely related. We therefore recommend users implement a rotating model comparison strategy when possible. It is important to note that the results from qpAdm are always going to depend on the availability of samples. Thus, even if the rotating strategy points to one particular model as the optimal model for a given dataset, this should not be taken as proof that the source populations identified are the actual best sources populations. For example, in Figure 1, if data were available from population 8 and not from population 9, the rotating model would identify populations 5 and 8 as the optimal sources of population 14. This would be correct, given the samples available, but it would come as no surprise if data from population 9 subsequently became available and it was deemed a better source than population 8. A number of examples exist in which previously identified qpAdm models have been refined when ancient DNA from new populations has become available, including in the Levant (Haber *et al.* 2017; Harney *et al.* 2018) and Sardinia (Haak *et al.* 2015; Chiang *et al.* 2018; Fernandes *et al.* 2020; Marcus *et al.* 2020).

While qpAdm’s ability to identify the optimal admixture model is affected by data quality, including the amount of missing data, the number of individuals in an analysis population, and the rate of ancient DNA damage, none of these factors ever bias qpAdm towards accepting a non-optimal model and rejecting the optimal model. Instead, we find that high rates of missing data or small sample size may make it more likely for qpAdm to accept multiple models. On the other hand, ancient DNA damage appears to cause qpAdm to be too stringent when it occurs at differential rates in the target and optimal source populations, often rejecting models that should be considered optimal, and resulting in biased admixture proportion estimates. While these results show that improving data quality and carefully curating data prior to analysis should be a priority of qpAdm users, they are promising as they suggest that data quality issues are unlikely to causes users to infer an incorrect model of admixture using qpAdm.

Although we find that the performance of qpAdm matches theoretical predictions under standard conditions, we also highlight several cases in which users should exercise caution. For instance, we find that users should attempt to limit the number of reference populations included in a qpAdm model, as the inclusion of too many reference populations may result in lowered p-values. Further, we show that qpAdm may produce plausible admixture proportion estimates and p-values in cases where the population of interest was not formed via admixture, such as the case of continuous migration, therefore users should be careful to consider whether alternative demographic models may better explain their data.

Overall, we find that qpAdm is a useful tool for identifying plausible admixture models and estimating admixture proportions, and that its performance matches theoretical expectations. qpAdm is particularly useful because it can be used in cases where the underlying population history of all the populations included in the analysis is difficult to determine and can therefore be used in cases where it may not be possible to use other tools for modeling population histories that involve admixture, like qpGraph and TreeMix. We include an updated user guide for qpAdm in Supplementary Materials 1 in order to make this method more accessible to future users.

## Notes

### Competing Interest Statement

The authors have declared no competing interest.

