## Supplementary Materials for "Assessing the Performance of qpAdm: A Statistical Tool for Studying Population Admixture"

Table of Contents

Figure S1. Distribution of p-values generated for various qpAdm models....................................2
qpAdm User Guide ........................................................................................................................3

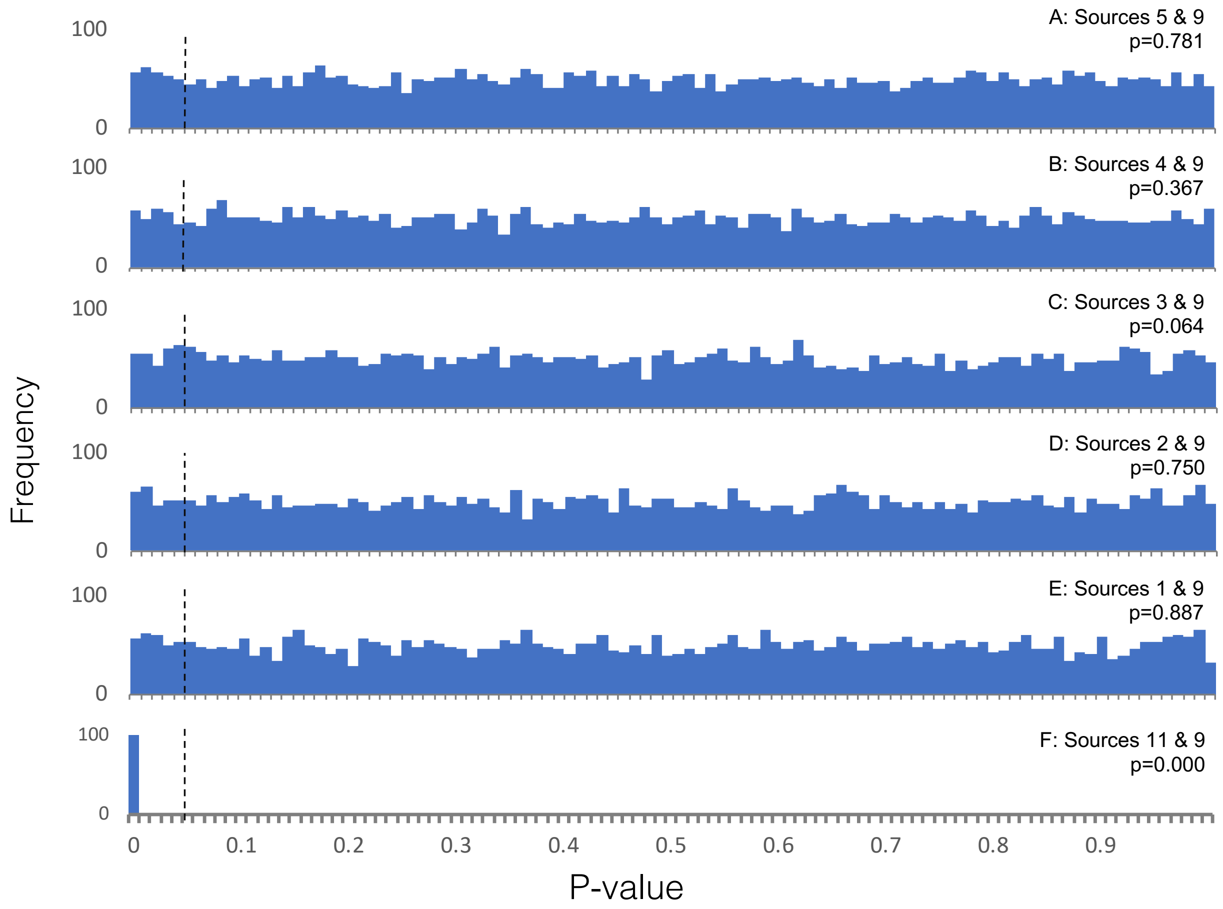

Figure S1. Distribution of p-values generated for various qpAdm models.

The distribution of p-values generated by 5,000 replicates of qpAdm is shown for all models, except when sources 11 & 9 are used, in which case only 100 replicates were generated. Panel A-F shows the distribution of p-values produced by models using population 9 as a source, in combination with population 1-5 or 11, respectively. As population 0 is not included as a reference population, models using sources 1-5 all serve as appropriate source populations for the ancestry of population 14. Vertical black dotted lines indicate the p-value threshold of 0.05, above which qpAdm models are considered plausible. The results of a Kolmogorov-Smirnov test to determine whether the p-values are uniformly distributed are indicated.

### qpAdm User Guide

#### Overview

qpAdm is a statistical tool for studying the ancestry of populations with histories that involve admixture between two or more source populations. Using qpAdm, users can assess the plausibility of admixture models and estimate admixture proportions.

qpAdm is written in the language C and can be downloaded on github as part of the AdmixTools package: https://github.com/DReichLab/AdmixTools

### Installation

#### Dependencies

qpAdm is part of the AdmixTools package. AdmixTools requires users to link copies of the following tools: GNU Scientific library (gsl), openblas, gfortran, and lapack. In order to use other versions of BLAS, users should update the Makefile with the corresponding version of BLAS.

For users building AdmixTools on a Mac*, the required dependencies can be installed with homebrew using the following commands:

```
brew install gsl
brew install openblas
```

*Users installing AdmixTools on a Mac must also uncomment the lines in the AdmixTools src/Makefile that modify the CFLAGS and LDFLAGS before installing AdmixTools. These parameters may need to be adjusted depending on the user's compute environment set up.

#### Download

qpAdm can be downloaded from github as part of the AdmixTools package (https://github.com/DReichLab/AdmixTools). To clone the AdmixTools github repository, use the following commands:

```
git clone https://github.com/DReichLab/AdmixTools.git
```

#### Compiling

All source code and executables for AdmixTools packages, including qpAdm, can be found in the src/ directory. To recompile the program, enter the AdmixTools directory and type:

```
cd src
make clobber
make all
make install
```

AdmixTools executables, including qpAdm, will be in ../bin

### Getting Started

#### Input Data

qpAdm can be run on data in the following 5 formats, which are supported by AdmixTools:

ANCESTRYMAP
EIGENSTRAT
PED
PACKEDPED
PACKEDANCESTRYMAP

For the fastest analyses, we recommend PACKEDANCESTRYMAP format. For full descriptions of each of these formats, see

https://github.com/DReichLab/AdmixTools/tree/master/convertf/README

##### Left Population File

The target and source populations are defined in this file. The first population included in the list is considered to be the target populations, and all other populations are considered to be potential sources of the ancestry in the target population. One population should be listed per line. The order of source populations (i.e. all populations after the first population) does not matter.

##### Right Population File

This is a list of reference populations to be included in the qpAdm model. The number of reference populations must be greater than the number of left (i.e. target and source) populations. One population should be listed per line. The first population in the list will be used as a base for all $f_4$-statistics calculated. Population order after the first population does not matter.

##### Parameter files

In order to run qpAdm, users must provide a parameter file (i.e. a "parfile") that contains pointers to the data and population model to be analyzed and indicates additional parameters to be used. The following parameters must be specified:

*Required parameters:*

genotypename: pointer to the input genotype file
snpname: pointer to the input snp file, corresponding to the defined genotype file
indivname: pointer to the input ind file, corresponding to the defined genotype file
popleft: pointer to the left population file (described above)
popright: pointer to the right population file (described above)

*Optional parameters include:*

| | |
|---|---|
| details: | Provides information about the difference between the fitted model and real data. See the output section for more information about the information that this option provides.<br>Default: YES |
| allsnps: | Specifies whether $f_4$-statistics will use the intersection of SNPs covered by all populations included in the model, or only the intersection of the 4 populations included in each $f_4$-statistic<br>Default: NO – restricts analysis SNP set to intersection of all SNPs among all populations<br>Alternative: YES – uses all available SNPs for each $f_4$-statistic comparison |
| chrom: | Specifies a single chromosome to be used in analysis<br>Default: NULL |
| nochrom: | Specifies a single chromosome to ignore during analysis. May be useful for a crude chromosomal jackknife<br>Default: NULL |
| numchrom: | Specifies the total number of chromosomes to be used in analysis. If “numchrom: 1” only chromosome 1 will be used, while if “numchrom: 22” all human autosomes will be used. It is recommended to set this number equal to the total number of autosomes in the organism being studied.<br>Default: 22 |
| diagplus: | By default, a constant is added along the diagonal of various matrices in order to make qpAdm output more robust results in boundary cases where the mixing coefficients are not well determined. In order to override this feature, set “diagplus: 0”<br>Default: NULL |
| hires: | Increases the number of decimal places reported for admixture proportions (in the “best coefficients” line) and standard errors (in the “std. errors” line) from 3 to 9 when set to “YES”<br>Default: NO |

| | |
|---|---|
| instem: | Allows users to specify a common prefix that is shared between all input data files, rather than defining each separately. For example, if the "instem: test" parameter is defined, qpAdm will expect the following input files: test.ind, test.snp, and test.geno<br>Default: NULL |
| hiprec_covar: | Prints error covariance matrix in high precision when set to YES<br>Default is to report the error covariance matrix multiplied by 1 million, high precision mode multiples by 1 billion<br>Default: NO |
| badsnpname: | Specifies a list of SNPs that are ignored during analysis. Each SNP should be listed on a single line and should be referred to by name (i.e. the first column in an EIGENSTRAT .snp file)<br>Default: NULL |
| blockname: | Allows users to specify custom block numbers for the block jackknife calculations. Specifies a list of SNPs to be analyzed. One SNP per line, followed by the desired block number (an integer greater than or equal to 1). SNPs assigned a block number of "-1" or that are excluded from the list will be ignored.<br>Default: NULL |
| blgsize: | The jackknife block size (in Morgans). Note qpAdm checks to make sure a reasonable number has been suggested here. If a block size is accidentally specified in centimorgans, this may be flagged and corrected by qpAdm during analysis.<br>Default: 0.05 |
| gfromp: | When this option is selected, the genetic distance defined in the snp input file is ignored, and qpAdm instead uses the physical distance as a proxy for genetic distance, assuming 100 million bases corresponds to 1 Morgan.<br>Default: NO |
| fancyf4: | When this option is selected, during $f_4$-statistic calculation, if statistics of the form $f_{4(}$A,B; C,D) are being calculated and if genotype information for population D is missing, in cases where A=B, the $f_4$-statistic is still considered, as it will always be equal to 0.<br>Default: YES |
| seed: | Specifies the seed to be used. If set to 0, a random seed will be chosen according to the system clock.<br>Default: 0 |

#### Running qpAdm

To run qpAdm, use the following command:

```
DIR/bin/qpAdm -p parfile
```

Where parfile is a pointer to the parameter file you have prepared for the analysis, and DIR is the path to the bin directory where qpAdm is stored. Users may optionally write results to a logfile (recommended).

#### Output

Below is an example of a typical qpAdm output. Annotations describing each section are preceded by '####' and highlighted in yellow.

```
#### A pointer to the parameter file used for analysis
/home/np29/o2bin/qpAdm: parameter file: qpAdm_v1_left_14_5_9_right_13_12_10_7_0_0.50_ds10000.par

### THE INPUT PARAMETERS
##PARAMETER NAME: VALUE

#### A copy of the parameter file used for analysis.

genotypename: scenario2_v1_a0.50_ds10000.geno
snpname: scenario2_v1_a0.50_ds10000.snp
indivname: scenario2_v1_a0.50_ds10000.ind
popleft: left_14_5_9
popright: right_13_12_10_7_0
allsnps: YES
details: YES
summary: YES

## qpAdm version: 1010        #### The version of qpAdm used
seed: 1164929463              #### The seed used for analysis. qpAdm chooses a random seed (using the clock) by default, but this can be set using the "seed"
optional parameter

#### Any errors or potential issues may be flagged here

#### A list of left populations used for analysis. The first population is the target population, all other subsequent populations serve as sources
left pops:
Pop_14
Pop_5
Pop_9

#### A list of right populations used as references in the analysis. The first population is used as a base for all f4-statistic calculations
right pops:
Pop_13
Pop_12
Pop_10
Pop_7
Pop_0
```

```
#### The number of individuals per population used in the analysis. Column 1- ordered list, Column 2- population ID, Column 3- # individuals per population
 0      Pop_14  10
 1      Pop_5   10
 2      Pop_9   10
 3      Pop_13  10
 4      Pop_12  10
 5      Pop_10  10
 6      Pop_7   10
 7      Pop_0   10

jackknife block size:    0.050       #### Size of the block jackknife (Default 0.050, can be set using the "blgsize" parameter)
snps: 10000  indivs: 80              #### Total number of SNPs in the dataset (not the total number of snps analyzed), Total number of individuals analyzed
number of blocks for block jackknife: 428     #### Total number of blocks used for block jackknife
## ncols: 10000                               #### Number of SNPs in dataset

#### The number of sites where at least one individual has coverage for each population. Column 1- Population name, Column 2- Number of sites
coverage:      Pop_14  10000
coverage:      Pop_5  10000
coverage:      Pop_9  10000
coverage:      Pop_13  10000
coverage:      Pop_12  10000
coverage:      Pop_10  10000
coverage:      Pop_7  10000
coverage:      Pop_0  10000
dof (jackknife):   346.407                    #### Effective number of blocks used in block jackknife
numsnps used: 10000                           #### Total number of SNPs analyzed
codimension 1                                 #### This line always reads codimension 1 for all qpAdm analyses

#### This section reports similar information as provided by the qpWave methodology, testing whether a matrix of maximum rank minus 1 (in this case 1) can be accepted

f4info:
#### f4 rank – the rank being tested
#### dof – the number of degrees of freedom in the analysis
#### chisq & tail – chi square and p values calculated from the matrix of f4-statistics
#### chisqdiff & taildiff – comparisons of the chisq and p-values associated with the rank under consideration versus that rank minus 1.

f4rank: 1 dof:     3 chisq:    14.028 tail:       0.00286708986 dofdiff:     5 chisqdiff:   -14.028 taildiff:        1
```

```
#### qpAdm calculates two matrices, matrix A is of size (# of left pops x rank) and B is of size (rank x # of right pops). These matrices are reported below. Each column should be multiplied by the corresponding scale value listed above it.

B:
      scale     1.000
        Pop_12    0.421
        Pop_10   -0.037
         Pop_7    0.937
         Pop_0    1.716
A:
      scale  2279.353
         Pop_5    0.588
         Pop_9   -1.286

#### Next, qpAdm considers whether a matrix of full rank (in this case rank=2) can be accepted. This section should be interpreted in the same way as the above section, unless otherwise noted

full rank
f4info:

##### taildiff compares the difference between p-values produced for the full rank versus full rank minus 1. This is the p-value reported by qpAdm. If this value is very small, the qpAdm model is likely incorrect

f4rank: 2 dof:     0 chisq:    0.000 tail:          1 dofdiff:     3 chisqdiff:    14.028 taildiff:      0.00286708986

B:
      scale  3702.746  1434.652
        Pop_12   -1.213   -0.782
        Pop_10   -0.304   -0.030
         Pop_7   -0.507   -1.143
         Pop_0    1.476   -1.443
A:
      scale     1.414     1.414
         Pop_5    1.414    0.000
         Pop_9    0.000    1.414
```

```
#### The estimated admixture proportions, order corresponds to that of left population list
best coefficients:     0.686     0.314
#### Mean admixture proportions computed by the block jackknife analysis.
#### Note if the jackknife mean and best coefficients estimates are very different, there is likely to be an issue (i.e. bizarre data in a few blocks)
Jackknife mean:      0.676547472     0.323452528
#### The estimated standard errors assigned to each admixture proportion
std. errors:     0.118     0.118

#### An error covariance matrix that is computed with the block jackknife.

error covariance (* 1,000,000)
     13820     -13820
    -13820      13820

#### An optional line produced using the "summary: YES" parameter. It reports
#### "summ: [target pop] [rank]       [p-value]          [admix prop 1] [admix prop 2] [error covariance] [error covariance] [error covariance]"

summ: Pop_14    2     0.002867     0.677     0.323     13820     -13820     13820

#### This section reports the qpAdm results that would be produced if the admixture estimate for one or more source populations is forced to be equal to 0
#### The "fixed pat" parameter (Column 1) indicates which populations are forced to be equal to 0 (0=not forced, 1 = forced)
#### The "wt" parameter (Column 2) reports the number of populations that are forced to have admixture proportion estimates equal to 0
#### The remaining columns report Column 3- degrees of freedom, Column 4- chi squared value, Column 5-tail probability, Column 6 & 7- assigned admixture proportions

    fixed pat  wt  dof    chisq       tail prob
          00  0     3   14.028     0.00286709     0.686     0.314          #### In this row, all source populations are used
          01  1     4   20.479     0.000401644     1.000     0.000         #### In this row, only the first source population (i.e. Pop_5 is used)
          10  1     4   52.554     1.05636e-10     0.000     1.000         #### In this row, only the second source population (i.e. Pop_9 is used)

#### The best pat section compares the tail prob when no pops are dropped from analysis with the highest tail prob produced when one pop is dropped
#### A p-value greater than 0.05 for the nested model suggests that it may be appropriate to drop one or more populations from the model

best pat:          00      0.00286709              - -
best pat:          01     0.000401644  chi(nested):     6.451 p-value for nested model:      0.0110916
```

```
####The following section is produced when the "details: YES" option is selected. It reports the difference between fitted model and real data. See the main text for an explanation of how to interpret this section

coeffs:     0.686     0.314

## dscore:: f_4(Base, Fit, Rbase, right2)
## genstat:: f_4(Base, Fit, right1, right2)

details:                Pop_5                Pop_12    -0.000328   -1.756199
details:                Pop_9                Pop_12    -0.000545   -2.898009
dscore:                Pop_12 f4:    -0.000396 Z:     -2.528301

details:                Pop_5                Pop_10    -0.000082   -0.402814
details:                Pop_9                Pop_10    -0.000021   -0.115325
dscore:                Pop_10 f4:    -0.000063 Z:     -0.379765

details:                Pop_5                 Pop_7    -0.000137   -0.821139
details:                Pop_9                 Pop_7    -0.000796   -4.840316
dscore:                Pop_7 f4:    -0.000344 Z:     -2.518733

details:                Pop_5                 Pop_0     0.000399    2.293766
details:                Pop_9                 Pop_0    -0.001006   -6.156302
dscore:                Pop_0 f4:    -0.000042 Z:     -0.296865

gendstat:               Pop_13                Pop_12    -2.528
gendstat:               Pop_13                Pop_10    -0.380
gendstat:               Pop_13                 Pop_7    -2.519
gendstat:               Pop_13                 Pop_0    -0.297
gendstat:               Pop_12                Pop_10     1.718
gendstat:               Pop_12                 Pop_7     0.309
gendstat:               Pop_12                 Pop_0     1.974
gendstat:               Pop_10                 Pop_7    -1.726
gendstat:               Pop_10                 Pop_0     0.132
gendstat:                Pop_7                 Pop_0     2.757

##end of qpAdm:       0.540 seconds cpu        9.150 Mbytes in use
```

#### Description of “details: YES” output

The optional parameter “details: YES” creates a section at the end of the qpAdm log file that describes the difference between the fitted model and real data. This comparison is reported in two ways, referred to as dscore and gendstat. Both parameters highlight the difference between the real target population (i.e. the “Base” population) and the modeled population that is produced by a weighted combination of the source populations (i.e. the “Fit” population).

##### *dscore:*

In the case of dscore, the difference between the real target population and this theoretical “Fit” population is calculated using $f_4$-statistics of the form $f_4$(Base, Fit; Rbase, right2) where *Rbase* is the primary reference population (i.e. the first population listed in the Right population list) and *right2* is all other populations in the right population list.

For each *right2* population, the results of this $f_4$-statistic is reported in the “dscore” section. In each case, the line reads:

“dscore: *right2* f4: [calculated f4-statistic] Z: [calculated z-score]”

Additionally, for each source population, there is a corresponding line labeled “details” above this dscore section, where the results of the $f_4$-statistic of the form $f_4$(Base, source; Rbase, right2) are reported separately, in the form:

“details: *source right2* [calculated f4-statistic] [calculated z-score]”

##### *genstat:*

The genstat data reports similar information, but rather than using the primary reference population in all calculations, results for the statistic $f_4$(Base, Fit; right1, right2) is reported for all combinations of reference populations, right 1 and right2, are reported in the form:

“gendstat: *right1 right2* [calculated z-score]”

#### Example Analysis

The following are examples of the input files required for running qpAdm. They were used to analyze a replicate the 10,000 SNP downsampled data with alpha 0.50 shown in Figure 3A of the main text. These files and the corresponding example data are provided in Supplementary File 4.

##### Parameter file:

qpAdm_v1_left_14_5_9_right_13_12_10_7_0_0.50_ds10000.par

```
genotypename: scenario2_v1_a0.50_ds10000.geno
snpname: scenario2_v1_a0.50_ds10000.snp
indivname: scenario2_v1_a0.50_ds10000.ind
popleft: left_14_5_9
popright: right_13_12_10_7_0
allsnps: YES
details: YES
summary: YES
```

##### popleft file:

left_14_5_9

```
Pop_14
Pop_5
Pop_9
```

##### popright file:

right_13_12_10_7_0

```
Pop_13
Pop_12
Pop_10
Pop_7
Pop_0
```

#### Running the example:

The qpAdm output from this analysis is annotated in the example output shown above. To run, type the following commands from a directory that contains all the example files.

```
DIR/bin/qpAdm -p qpAdm_v1_left_14_5_9_right_13_12_10_7_0_0.50_ds10000.par
```

Where DIR is the path to the AdmixTools directory.

### Usage Recommendations

#### Data type

qpAdm supports all data formats supported by the AdmixTools package (EIGENSTRAT, ANCESTRYMAP, PED, PACKEDPED, and PACKEDANCESTRYMAP). In order to increase the speed of analysis, we recommend using data in the PACKEDANCESTRYMAP format whenever possible.

#### Parameters

In addition to the required parameters, we recommend using the following optional parameters:

| | |
|---|---|
| details: YES | This will provide additional output information that highlights the difference between the actual target population and the model fitted by qpAdm. See the *Description of "details: YES" output* section for further information. |
| summary: YES | This option will provide an easy to search summary line (labeled "summ:") in the output that includes the assigned p-value and admixture proportions for the proposed model. |
| allsnps: YES | The "allsnps: YES" option was developed in order to increase the number of SNPs analyzed by qpAdm in cases where very little SNP overlap exists between all populations included in the model. Rather than only analyzing sites that have available data for all populations included in the model, with the "allsnps: YES" option specified, each $f_4$-statistic is calculated using the SNP sites that are shared between the four populations involved in each statistic. While the choosing "allsnps: YES" option violates the underlying theoretical expectations of qpAdm, in practice, this option appears to improve qpAdm's ability to distinguish between optimal and non-optimal models and provides more accurate admixture proportion estimates when analyzing data with high rates of missing data. |

#### Selecting Left and Right Populations

When selecting populations to include in a qpAdm model, users should try to maximize the data quality by choosing populations that contain a large number of individuals (if they can be confidently grouped) or that contain individuals with as high coverage as possible.

##### Recent Gene Flow

qpAdm assumes that there has been no gene flow between the left and right populations following the admixture event of interest. Therefore, users should try to avoid including populations that are known to have experienced recent gene flow with one another whenever possible.

##### Ancient DNA damage

We recommend that users avoid analyzing qpAdm models that contain mixtures of modern and ancient populations in either the left or right set, as such mixtures can cause target or source populations to produce an artifactual signal of shared drift with reference right populations. Even within a set of target and source populations that are ancient, users should also be wary of scenarios where there are variable rates of ancient DNA damage across samples when analyzing datasets including transition polymorphisms that are susceptible to such damage. As our simulations have shown, this scenario can cause admixture proportion estimates produced by qpAdm to be biased away from the true admixture proportion as samples with similar damage rates can appear artifactually too closely related to each other; in such a scenario, it is best to restrict to transversion polymorphisms not vulnerable to ancient DNA damage.

##### The first right population

In order to simplify calculations, qpAdm uses the first population that is specified in the right population list as a base for all $f_4$-statistics that it calculates. It is therefore important to choose this population with care.

If the “allsnps: YES” option is selected, be sure to select a population that is of relatively high coverage to include in this position, as all $f_4$-statistics calculated will still be restricted to only using sites that are covered in this individual.

Additionally, as slight variations in the results produced by qpAdm when different reference populations are placed in this first position are expected, it is recommended that the same reference population be specified in this position across all models that are being compared, whenever possible. Common practice is to specify a population that is unlikely to be closely related to the admixture event being considered, so as not to create significant biases in the analyses if it has to be removed from the list of right populations to be used as a source population in comparison analyses.

#### Choosing informative right populations

qpAdm harnesses differential relatedness between left and right populations in order to determine whether a particular model is plausible and to assign admixture proportions. Only cases where a reference population is differentially related to the target and source populations—resulting in differences in the value of the statistics produced for each of the left populations—are informative for determining whether the model is plausible and for calculating admixture proportions. Therefore, the inclusion of a right (or reference) population that is symmetrically related to all left populations does not add meaningfully to the qpAdm model.

In cases where all right populations are symmetrically related to all left populations, qpAdm will likely be unable to reject any models, and will report arbitrary admixture proportion estimates with very high standard errors. One way to avoid this issue is to prescreen the right populations included in the qpAdm model, to be sure that they are differentially related to the left populations in at least some proportion of the $f_4$-statistics that will be calculated. To do this, we recommend running $f_4$-statistics of the form $f_4(Left_i, Left_j; Right_k, Right_l)$ for all combinations of *Left* and *Right* populations prior to analysis. If a *Right* population never produces significant $f_4$-statistics, it is not an informative reference population.

In some cases it may be useful to include some number of right populations that are not differentially related to the left populations included in the model. For instance, when trying to compare a variety of models that use different left populations, it may be preferable to use the same set of right populations, even in cases where some right populations are only informative for some of the models being tested.

#### Optimal number of right populations

We recommend minimizing the number of right populations included in a model whenever possible, as when the number of right populations is large, the covariance matrix of $f_4$-statistics is likely to be poorly estimated. In the main text, we show that in cases where a very large number of right populations are included in a model, the p-values calculated by qpAdm are strongly biased towards 0. This effect is likely to result in the rejection of plausible models. We observe this effect when as few as 35 right populations are included in the model, however we caution that the number of right populations that can be safely included in a qpAdm model is likely to be highly dependent on data quality and the relative relationships of the populations included in the model to one another. We therefore caution users to be cognizant of this potential effect and limit the number of populations that are included in the right population set.

#### Comparing qpAdm Models

One of the primary objectives of qpAdm users is to identify an optimal model of the ancestry of a target population out of a variety of possible models. While there are a number of valid approaches for identifying this optimal model, there are many factors to consider when choosing a strategy for comparing possible models, including

1) Ensuring that the model includes right populations that are differentially related to the various source populations that are being used as potential left populations.
2) Ensuring that models are directly comparable. It is not appropriate to compare two models that use entirely different sets of right populations. While it may not be possible to use identical sets of right populations for all models under consideration, the right population sets should be as similar as possible.

While many strategies have been implemented by qpAdm users to directly compare models, we recommend using a “rotating” population approach, in which a set of populations of interest are selected to act either as source or reference populations. Users can then create models in which all possible combinations of source populations are defined in the left population list, and all remaining populations in the set that are not defined as source populations are set as right populations, to act as references. Based on simulated data, this strategy appears to maximize qpAdm’s ability to distinguish between possible sources of ancestry.

Users should note that this strategy is optimal in cases where there are a limited number of possible source populations to consider. In cases where users wish to consider a very large number of possible sources, it may be optimal to instead choose a smaller number of populations to act as right populations for all models being considered, in order to avoid producing qpAdm models with reduced p-values due to the inclusion of an excessive number of right populations (see the “Optimal number of right populations” section).

### About qpAdm

#### Citing qpAdm

qpAdm was first introduced in Haak et al (2015) and is described in Supplementary Materials Section 10.

An MLA version of this citation is provided below:

Haak, Wolfgang, et al. "Massive migration from the steppe was a source for Indo-European languages in Europe." *Nature* 522.7555 (2015): 207.

A bibtex version of this citation is also provided:

```
@article{haak2015massive,
  title={Massive migration from the steppe was a source for Indo-European
languages in Europe},
  author={Haak, Wolfgang and Lazaridis, Iosif and Patterson, Nick and Rohland,
Nadin and Mallick, Swapan and Llamas, Bastien and Brandt, Guido and Nordenfelt,
Susanne and Harney, Eadaoin and Stewardson, Kristin and others},
  journal={Nature},
  volume={522},
  number={7555},
  pages={207},
  year={2015},
  publisher={Nature Publishing Group}
}
```

#### Contact

Nick Patterson

#### Software Copyright Notice Agreement

This software and its documentation are copyright (2010) by Harvard University and The Broad Institute. All rights are reserved. This software is supplied without any warranty or guaranteed support whatsoever. Neither Harvard University nor The Broad Institute can be responsible for its use, misuse, or functionality. The software may be freely copied for non-commercial purposes, provided this copyright notice is retained.
