## Supplementary File 4 for "Assessing the Performance of qpAdm: A Statistical Tool for Studying Population Admixture": README

To run this qpAdm example, type the following commands from within the directory that contains the example files, replacing "DIR" with a pointer to the AdmixTools directory

DIR/bin/qpAdm -p qpAdm_v1_left_14_5_9_right_13_12_10_7_0_0.50_ds10000.par
